# Single-atom inhibition of oncogenic drivers through cysteine coordination

**DOI:** 10.64898/2026.08.27.747489

**Authors:** Wencong Zhao, Zhen Chen, Kai Cao, Wendi Huo, Yunjiang Zhang, Sixu Chen, Dongfang Xia, Qing Yuan, Peng Cao, Shaorui Sun, Xueyun Gao

**Author notes:** These authors contributed equally to this work (Wencong Zhao, Zhen Chen, Kai Cao).

## Abstract

Small-molecule inhibitors rely on molecular recognition within suitable binding pockets, leaving many disease-associated proteins difficult to target. Here, we introduce the concept of a single-atom inhibitor in which gold (Au) engages critical cysteine residues of oncogenic drivers to suppress their activity. We used an AI-assisted few-shot learning approach to identify EGFR-targeting peptides for *in vivo* Au delivery and showed that the lead candidate, 10714, promoted Au accumulation in EGFR-expressing cells and tumors. *In vivo*, Au exploited its intrinsic affinity for cysteine to inhibit two structurally distinct oncogenic proteins, engaging Cys797 in EGFR T790M and the mutation-derived Cys12 in KRAS G12C adjacent to their respective nucleotide-binding pockets. Structural and computational analyses supported stabilization of inactive nucleotide-bound states, while mutation of these cysteine residues abrogated Au-mediated inhibition. 10714-Au consequently suppressed oncogenic signaling, reduced non-small-cell lung cancer cell viability, and inhibited tumor growth in EGFR- and KRAS-mutant xenograft models and patient-derived organoids. These findings establish proof of principle for single-atom inhibition across structurally distinct oncogenic drivers and suggest that localized atomic coordination could provide an alternative mode of target engagement to conventional pocket-dependent inhibition.

## Introduction

Over the past two decades, small-molecule inhibitors have transformed cancer therapy, achieving substantial clinical success across a range of malignancies[1, 2]. However, their development generally depends on the presence of a binding site with suitable structural and chemical properties to support productive molecular interactions[3]. Many oncogenic drivers therefore remain difficult to target, even when they contain functional pockets that bind endogenous ligands. Indeed, the druggability of nucleotide-binding pockets varies markedly among oncogenic drivers. Whereas the ATP-binding pocket of the receptor tyrosine kinase epidermal growth factor receptor (EGFR) has been successfully exploited by small-molecule inhibitors[4], direct targeting of the nucleotide-binding pocket of the small GTPase KRAS has proved particularly challenging owing to its high affinity for guanine nucleotides and the absence of a readily exploitable binding environment[5]. These limitations motivate alternative strategies for inhibiting oncogenic drivers that are less constrained by pocket druggability.

An alternative approach would be to develop an inhibitor sufficiently small to access spatially restricted binding environments, with an intrinsic capacity for covalent engagement[6]. Reducing such an inhibitor to the atomic scale could, in principle, lessen its dependence on the depth, shape or surface chemistry of the surrounding pocket, while covalent interaction with a critical residue could provide stable target engagement. Together, these properties could enable inhibition of targets that are poorly amenable to conventional pocket-based drug design.

Metal ions offer one potential means of achieving such atomic-scale inhibition because they can regulate protein activity through coordination interactions[7, 8]. Gold offers an attractive candidate for this approach because of its capacity to form stable interactions with thiolate groups. In previous work, we found that a single gold atom (Au) potently inhibited human thioredoxin reductase through covalent Au–S engagement of the thiolate group of Cys189[9]. This finding raised the possibility that Au-thiolate chemistry could be exploited to inhibit other proteins containing functionally important cysteine residues. Notably, cysteine residues occur within or adjacent to the ATP- or GTP-binding pockets of several oncogenic drivers[10, 11], and cysteine residues are also frequently acquired through oncogenic mutations[12, 13]. We therefore hypothesized that single-atom Au–cysteine engagement could provide a means of inhibiting oncogenic drivers without requiring an extensively druggable binding pocket.

Here, we tested this concept using EGFR and KRAS as clinically important but structurally distinct oncogenic drivers in non-small cell lung cancer (NSCLC). To enable tumor-targeted delivery of gold, we developed a few-shot learning approach to generate an EGFR-targeting peptide. Using structural, biochemical and preclinical approaches, we show that single Au atoms can engage critical cysteine residues within or adjacent to the nucleotide-binding pockets of EGFR and KRAS, stabilizing their inactive states and suppressing oncogenic signaling. Together, these findings establish single-atom inhibition as an alternative framework for targeting oncogenic drivers with distinct pocket properties.

## Results

### Few-shot learning identifies candidate EGFR-targeting peptides

To evaluate the potential of single Au atoms to regulate protein activity *in vivo*, an effective targeted delivery system is required, as systemic administration of free metals can be limited by instability, nonspecific distribution, off-target toxicity and insufficient tumor accumulation[14, 15]. We therefore focused on designing EGFR-targeting peptides for NSCLC cells, since EGFR is reported to be highly expressed in over 60% of NSCLC cases and undergoes receptor-mediated internalization, providing a potential route for targeted gold delivery[16–18].

Because only a limited number of experimentally validated EGFR-targeting peptides were available for model training, we developed a few-shot learning model based on a long short-term memory (LSTM) network (**Figure 1a**). We pre-trained a *de novo* peptide-generation model and subsequently fine-tuned it using 54 experimentally validated EGFR-targeting peptides to learn recurrent sequence motifs and residue-dependency patterns associated with EGFR recognition. We then coupled the learned sequence distribution with mutation-based sampling to explore a broader sequence space, generating 12,000 candidate peptides per batch that retained characteristics of the experimental training set (**Figure 1b, Table S1**). Principal component analysis (PCA) showed that the generated and experimental peptides occupied similar chemical space (**Figure S1a**). Similar PCA distributions were obtained across three independently generated batches (**Figure S1b**), indicating consistency in the chemical properties of the generated peptide populations.

**Figure 1.**
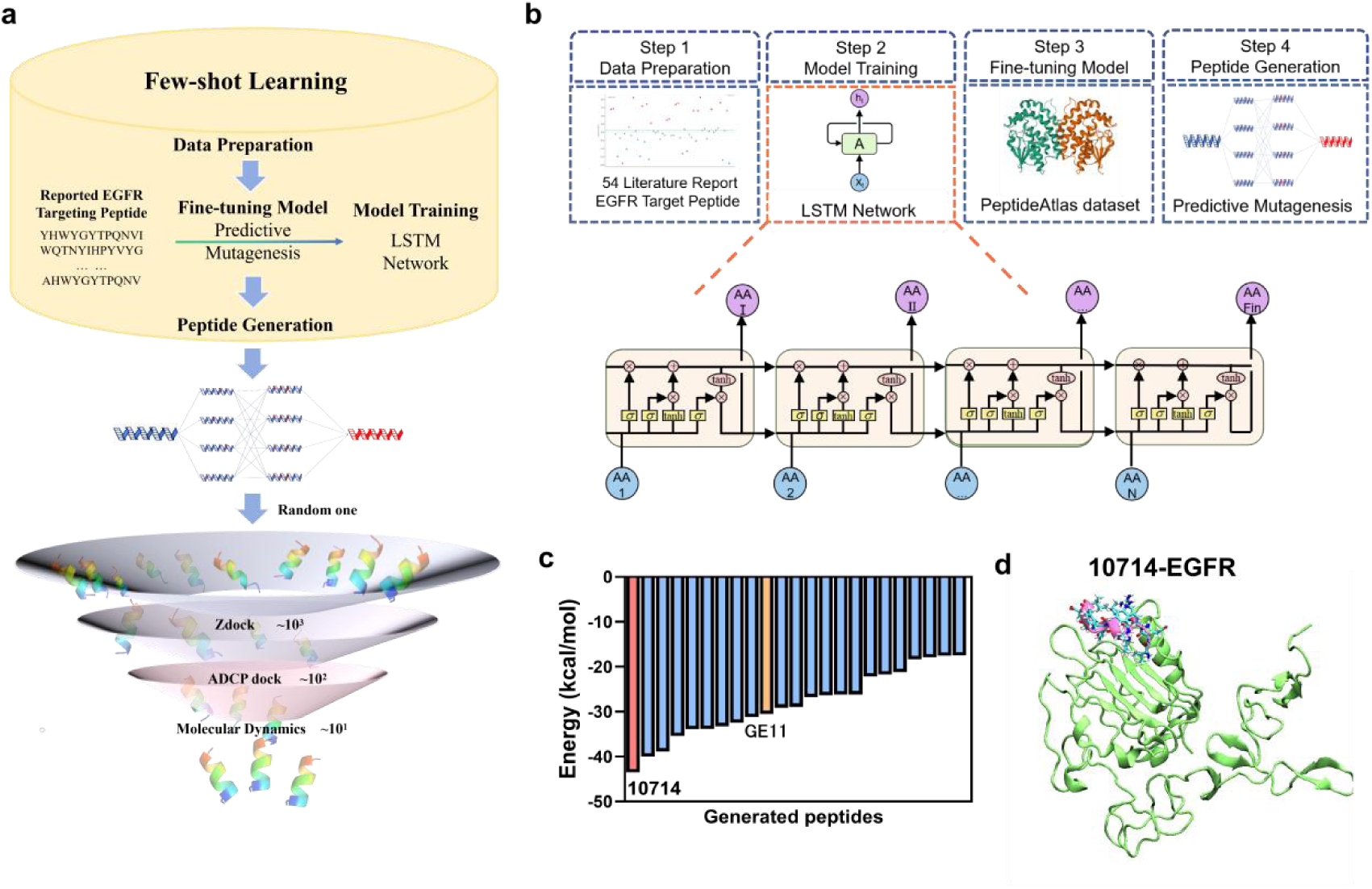
Few-shot learning identifies candidate EGFR-targeting peptides. **(a)** Workflow for the generation and computational screening of EGFR-targeting peptide candidates. **(b)** Architecture of the few-shot learning model used to generate EGFR-targeting peptide candidates. **(c)** Simulated non-bonded interaction energies between generated peptide candidates and the extracellular region of EGFR. Peptide 10714 and the experimentally validated EGFR-targeting peptide GE11 are highlighted in red and orange, respectively. **(d)** Molecular dynamics (MD) simulation of the predicted complex between peptide 10714 and the extracellular region of EGFR.

We next predicted three-dimensional structures for the generated and experimental peptides and performed molecular docking against EGFR. Using the highest ZDOCK score among the experimental peptides (1,549.433 for GE11[19]) as a benchmark, we identified 753 generated candidates that met this threshold for further screening using AutoDock CrankPep (ADCP) (**Tables S2, S3**). The 20 top-scoring peptide–EGFR complexes were subsequently subjected to molecular dynamics (MD) simulations, and their non-bonded interaction energies were calculated (**Figure 1c**, **Table S2**). This analysis identified five leading candidates (10714, 1648, 697, 4959 and 3469), which also showed favorable ZDOCK and ADCP scores (**Tables S3, S4**). Among these candidates, peptide 10714 showed the strongest predicted interaction with EGFR (**Figure 1d, Figure S2**).

### Characterization of peptide-Au molecules and their EGFR-targeting properties

To enable gold conjugation, we modified the peptides with a C-terminal YCC motif for Au–S coordination, separated from the targeting sequence by a hydrophilic KKKK spacer. Peptide 4959 showed poor aqueous solubility and was therefore excluded from subsequent experimental analyses. We conjugated the remaining four generated peptides and the previously reported GE11 peptide to gold using a one-step chemical method. All resulting peptide–Au molecules exhibited blue fluorescence (**Figure 2a, Figure S3a**), and UV–Vis absorption spectroscopy revealed a characteristic red shift relative to the corresponding unconjugated peptides, consistent with successful conjugation (**Figure 2b, Figure S3b**). Dynamic light scattering (DLS) and zeta-potential measurements showed narrow hydrodynamic size distributions and negative surface charges, respectively (**Figure 2c,d, Figure S3c,d)**. Matrix-assisted laser desorption/ionization time-of-flight mass spectrometry (MALDI-TOF-MS) supported a consistent Au₅Peptide₃ composition (**Figure 2e, Figure S3e**). Computational modelling and structural optimization further predicted coordination of the peptide thiol groups with the Au core through Au–S interactions (**Figure 2f**).

**Figure 2.**
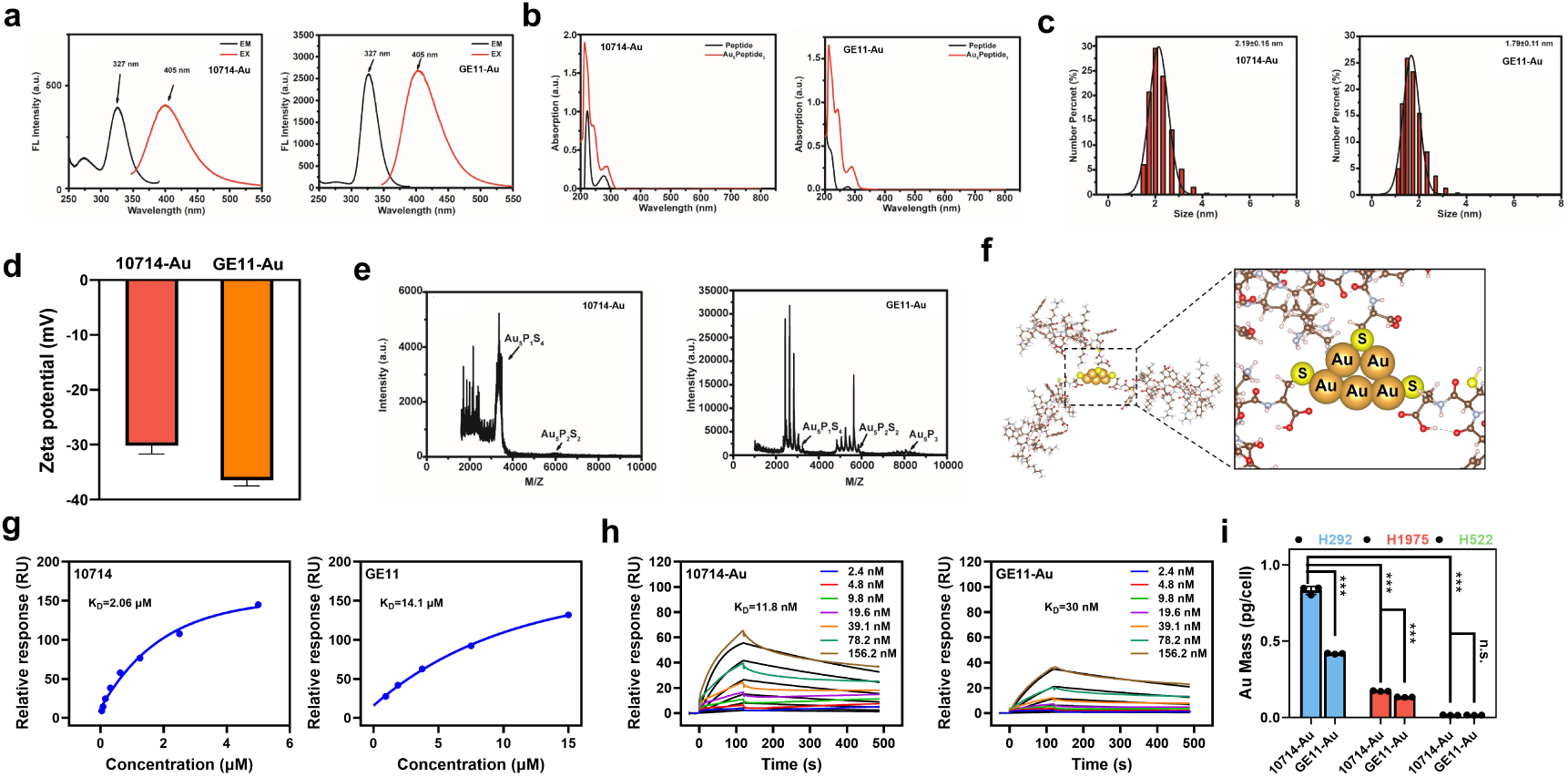
Characterization of 10714-Au and GE11-Au and their EGFR-targeting properties. **(a)** Fluorescence excitation (black) and emission (red) spectra of 10714–Au and GE11–Au. **(b)** UV–Vis absorption spectra of 10714–Au and GE11–Au compared with the corresponding unconjugated peptides. **(c)** Dynamic light scattering (DLS) analysis of 10714– Au and GE11–Au. **(d)** Zeta potentials of 10714–Au and GE11–Au. **(e)** MALDI-TOF-MS spectra of 10714–Au and GE11–Au acquired in positive-ion linear mode. The proposed composition of the peptide–Au molecules is Au_5_Peptide_3_. **(f)** Computationally optimized three-dimensional structure and predicted coordination mode of 10714–Au. The left panel shows the overall optimized conformation, and the enlarged view highlights the predicted coordination between gold atoms (Au, gold spheres) and peptide sulfur atoms (S, yellow spheres). **(g)** SPR analysis of the binding of peptides 10714 and GE11 to the extracellular domain of EGFR. **(h)** SPR analysis of the binding of 10714–Au and GE11-Au to the extracellular domain of EGFR. **(i)** Cellular Au accumulation following incubation of 10714– Au or GE11–Au with NSCLC cell lines with different levels of EGFR expression, measured by ICP-MS.

We next used surface plasmon resonance (SPR) to measure the EGFR-binding affinities of the EGFR-targeting peptides and corresponding peptide-Au molecules. Among the generated EGFR-targeting peptides, peptide 10714 showed the strongest binding, with a *K*_D_ of 2.41 μM compared with 6.09 μM for the experimentally validated peptide GE11[19] (**Figure 2g, Figure S4a**). Gold conjugation increased the apparent EGFR-binding affinities of all tested peptides, with peptide 10714-Au showing an approximately 200-fold decrease in *K*_D_ (11.8 nM; **Figure 2h, Figure S4b**).

To assess whether cellular gold uptake varied with EGFR expression, we measured Au accumulation by inductively coupled plasma mass spectrometry (ICP-MS) across three NSCLC cell lines with different EGFR expression levels (**Figure S5a**). All peptide-Au molecules showed the greatest Au accumulation in EGFR-high H292 cells and the lowest accumulation in EGFR-low H522 cells (**Figure 2i, Figure S5b**). Moreover, 10714–Au showed greater cellular Au accumulation than GE11–Au across all three cell lines (**Figure 2i**), supporting its selection as the lead EGFR-targeting delivery molecule.

### 10714-Au inhibits oncogenic EGFR and KRAS signaling and reduces NSCLC cell viability

Having established 10714–Au as the lead EGFR-targeting delivery molecule, we next asked whether the peptide–Au molecules could inhibit the activity of key oncogenic drivers in NSCLC. Activating EGFR mutations can drive oncogenic signaling in NSCLC, while EGFR L858R/T790M is associated with acquired resistance to earlier-generation EGFR tyrosine kinase inhibitors (TKIs)^21,22^. 10714– Au potently inhibited EGFR L858R/T790M enzymatic activities via an *in vitro* kinase assay using purified intracellular domain of the mutant protein, with an IC₅₀ of 28 nM compared with 793 nM for GE11–Au (**Figure 3a**). This greater inhibitory potency was associated with higher binding affinity of 10714–Au for mutant EGFR (**Figure S6a**). Both peptide–Au molecules also showed greater inhibition of mutant EGFR than wild-type EGFR, accompanied by higher binding affinity for the mutant protein (**Figure S6a,b**).

**Figure 3.**
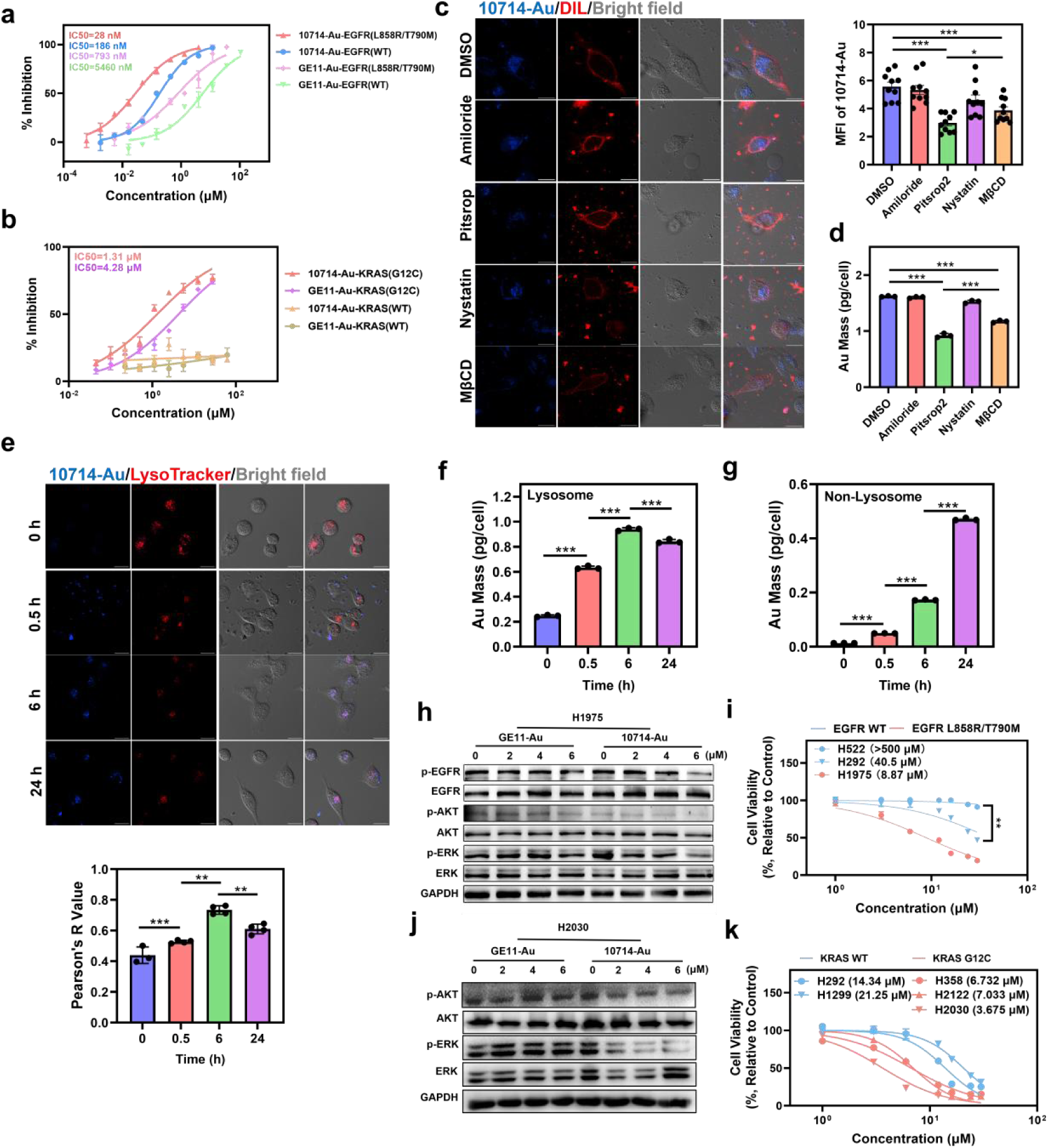
10714-Au inhibits oncogenic EGFR and KRAS signaling and reduces NSCLC cell viability. **(a,b)** Inhibition of the enzymatic activity of EGFR L858R/T790M (a) and KRAS G12C (b) by 10714–Au and GE11–Au. **(c)** H1975 cells were incubated with 10714– Au in the presence of inhibitors of different endocytic pathways and stained with DiI to visualize the cell membrane. Representative confocal images and quantification of intracellular 10714–Au fluorescence are shown. Scale bar, 20 μm. **(d)** Cellular Au accumulation in H1975 cells treated as in c, measured by ICP-MS. **(e)** H1975 cells were incubated with 10714–Au for the indicated times and stained with LysoTracker. Representative confocal images and Pearson correlation coefficients for colocalization are shown. Scale bar, 20 μm. **(f, g)** Au content in isolated lysosomal (f) and remaining non-lysosomal (g) fractions of H1975 cells following incubation with 10714–Au, measured by ICP-MS. **(h)** EGFR–AKT–ERK signaling in H1975 (EGFR L858R/T790M) cells following treatment with the indicated concentrations of 10714–Au or GE11–Au, assessed by western blotting. **(i)** Viability of NSCLC cell lines with different EGFR expression levels following treatment with the indicated concentrations of 10714–Au, measured using the CCK-8 assay. **(j)** AKT–ERK signaling in H2030 (KRAS G12C) cells following treatment with the indicated concentrations of 10714–Au or GE11–Au, assessed by western blotting. **(k)** Viability of NSCLC cell lines harboring wild-type (WT) or G12C-mutant KRAS following treatment with the indicated concentrations of 10714–Au, measured using the CCK-8 assay.

We then asked whether this inhibitory activity could extend to the structurally and functionally distinct small GTPase KRAS. Both molecules inhibited KRAS G12C, with 10714–Au again showing greater inhibitory potency and binding affinity than GE11–Au (**Figure 3b, Figure S7)**. By contrast, neither molecule inhibited wild-type KRAS, consistent with a requirement for the acquired Cys12 residue. Thus, 10714– Au inhibited two mechanistically distinct oncogenic drivers, with preferential activity towards their mutant forms.

Because both the EGFR kinase domain and KRAS are intracellular[20, 21], we next investigated 10714–Au cellular uptake and intracellular trafficking. Using its intrinsic blue fluorescence, we monitored 10714–Au accumulation in H1975 cells in the presence of inhibitors of different endocytic pathways. Inhibition of clathrin-mediated endocytosis substantially reduced intracellular 10714–Au fluorescence (**Figure 3c**), consistent with clathrin-mediated endocytosis as a major route of cellular uptake. ICP-MS analysis similarly showed reduced cellular Au accumulation following inhibition of clathrin-mediated endocytosis (**Figure 3d**). Following internalization, 10714–Au fluorescence initially colocalized with lysosomes and subsequently decreased over time (**Figure 3e**). Consistent with redistribution from lysosomes to the cytosol, ICP-MS analysis showed a time-dependent decrease in lysosomal Au accompanied by increased Au in the remaining non-lysosomal fraction (**Figure 3f, g**).

Having established cellular uptake of 10714–Au, we next examined its effects on downstream oncogenic signaling and cell viability. In EGFR-mutant H1975 cells, 10714–Au induced dose-dependent suppression of the EGFR–AKT–ERK signaling axis, with greater activity than GE11–Au (**Figure 3h**). Accordingly, 10714–Au showed dose-dependent cytotoxicity, with an IC₅₀ of 8.87 μM. This effect was greater in EGFR-mutant than EGFR-wild-type cells and in EGFR-high H292 than EGFR-low H522 cells (**Figure 3i**). Notably, the corresponding unconjugated peptides showed negligible cytotoxicity, indicating that Au conjugation was required for this activity (**Figure S8**). Consistent with these findings, both peptide–Au molecules preferentially induced apoptosis in EGFR-mutant cells, with 10714–Au showing greater activity (**Figure S9**).

In KRAS G12C H2030 cells, 10714–Au similarly suppressed AKT–ERK pro-survival signaling[20] (**Figure 3j**) and showed greater cytotoxicity in KRAS G12C than KRAS-wild-type cell lines (**Figure 3k, Figure S10**). Notably, 10714–Au also reduced the viability of KRAS-wild-type cells despite its lack of biochemical inhibition of wild-type KRAS. This effect could potentially reflect EGFR expression in these cells **(Figure S11a**). Moreover, cellular Au accumulation was lowest in EGFR-low H522 cells (**Figure S11b**), consistent with a contribution of EGFR expression to peptide–Au uptake. Collectively, these findings support a model in which EGFR-associated cellular uptake and intracellular trafficking of 10714–Au enable inhibition of mutant EGFR and KRAS, suppression of downstream pro-survival signaling and reduced NSCLC cell viability.

### Single-atom Au engages Cys797 of EGFR T790M and stabilizes an inactive state

Having established that 10714–Au inhibits mutant EGFR, we next investigated whether and how Au directly engages the EGFR kinase domain. We solved the 3.1 Å crystal structure of the EGFR T790M kinase domain (residues 696–1022) following incubation with 10714–Au (**Table S5**). The asymmetric unit contained two dimers comprising four EGFR kinase-domain molecules (**Figure S12a**). Both the *Fo*–*Fc* map and the anomalous difference Fourier map identified a single Au atom associated with each EGFR monomer (**Figure 4a, Figure S12b**). The Au atom was localized at Cys797, with an Au–S distance of 2.3 Å, consistent with direct Au–S coordination (**Figure 4b, Figure S12c**). The Au-binding environment also involved Asp800 and Arg841 and interactions with the phosphate group of AMP-PNP (non-hydrolyzable ATP analog), supporting stabilization of the EGFR complex in a nucleotide-bound inactive form (**Table S6-1**).

**Figure 4.**
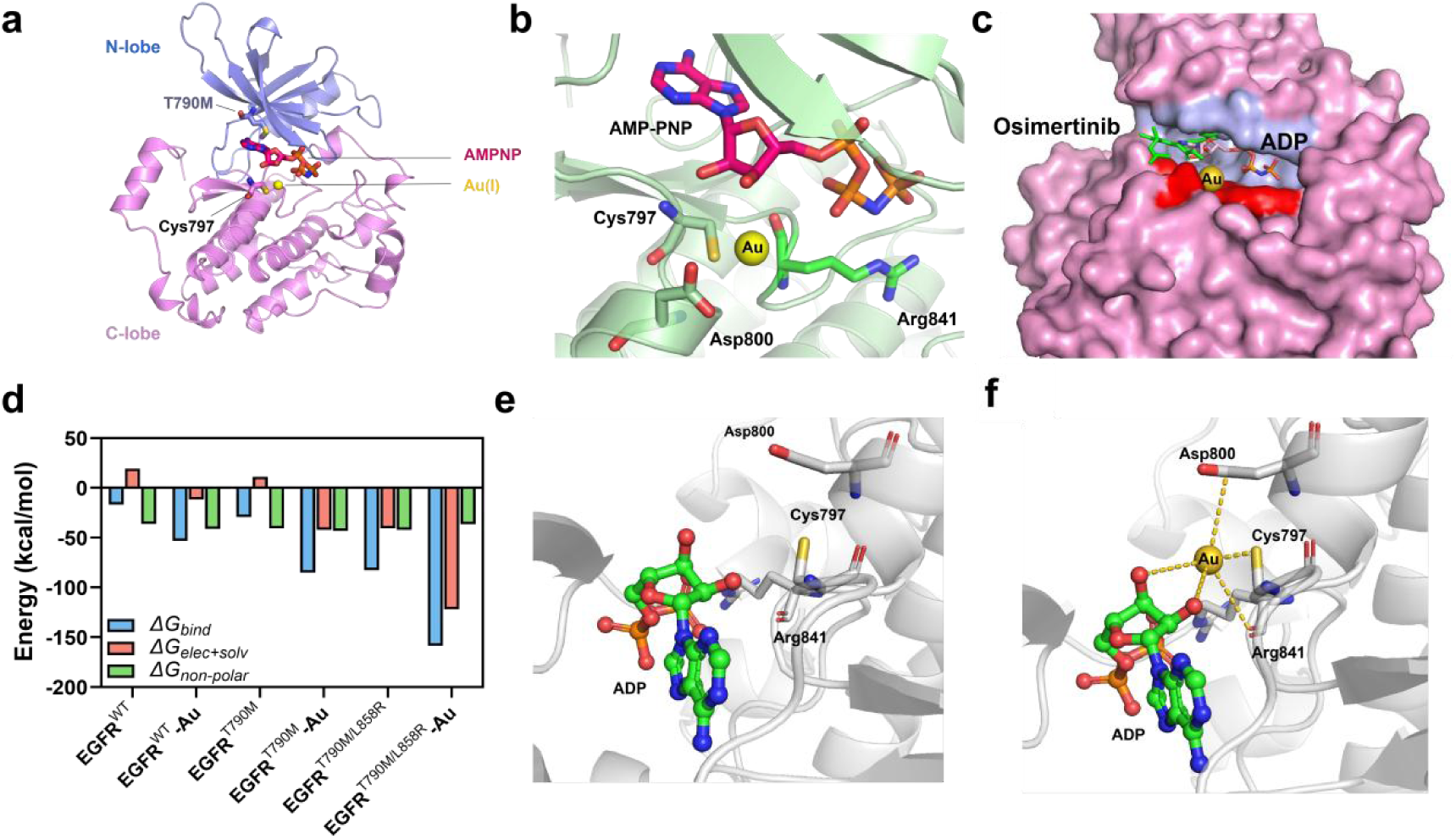
Single-atom Au engages Cys797 of EGFR T790M and stabilizes an inactive state. **(a)** Structure of one Au-bound EGFR T790M kinase-domain monomer (residues 696– 1022). The N-lobe is shown in light blue and the C-lobe in pink. AMP-PNP, Cys797 and T790M are shown as sticks. The Au–S interaction with Cys797 is indicated by a dashed line (distance, 2.3 Å). **(b)** Enlarged view of the Au coordination environment, showing AMP-PNP and the surrounding residues as sticks and the Au atom as a yellow sphere. **(c)** Superposition of the Au-bound EGFR structure with osimertinib-bound EGFR (PDB 6JX4). Osimertinib is shown as green sticks and occupies an extended region of the ATP-binding pocket (light blue), whereas Au is localized within a spatially confined coordination environment (red) around Cys797 at the edge of the nucleotide-binding pocket. **(d)** Predicted non-bonded interaction energy of ADP with wild-type EGFR, EGFR T790M or Au-bound EGFR T790M. **(e, f)** Predicted binding modes of ADP with EGFR T790M (e) and Au-bound EGFR T790M (f) from MD simulations. Predicted interactions involving Au are indicated by gold dashed lines and detailed in **Table S6-2**.

Interestingly, the Au-bound EGFR structure was highly similar to the untreated structure (PDB: 4ZSE), with conformational differences largely confined to loop regions (**Figure S12d**). By contrast, comparison with the structure of mutant EGFR bound to the covalent inhibitor osimertinib (PDB: 6JX4) showed that the Au-bound EGFR structure exhibited conformational differences in both the N- and C-lobes surrounding the ATP-binding pocket (**Figure S12e**). Whereas osimertinib occupies an extended region of the ATP-binding pocket, the Au atom was localized within a spatially confined coordination environment around Cys797 at the edge of the nucleotide-binding pocket, without occupying the entire pocket (**Figure 4c, Figure S12f, Table S6-2**). Thus, although both osimertinib and Au engage Cys797, the single Au atom engages EGFR through a highly localized coordination environment rather than the extended molecular interactions used by a small-molecule inhibitor.

MD simulations further suggested that Au binding increased the predicted non-bonded interaction between ADP and EGFR, stabilizing the ADP-bound complex and potentially restricting nucleotide exchange (**Figure 4d**). The simulations also predicted Au–S coordination with Cys797 together with four additional interactions involving the nucleotide-binding environment (**Figure 4e,f, Table S6-2**). Importantly, mutation of Cys797 to serine (C797S) abrogated the inhibitory effects of both 10714– Au and GE11–Au on EGFR (**Figure S12g**), providing functional support for a critical role of Cys797 in Au-mediated EGFR inhibition.

### Single-atom Au engages Cys12 of KRAS G12C and stabilizes an inactive state

Having established that single-atom Au engages Cys797 within the EGFR kinase domain, we next asked whether a similar mechanism could operate in the structurally distinct nucleotide-binding environment of KRAS G12C. We determined the 1.4 Å crystal structure of KRAS G12C following incubation with 10714–Au (**Table S5**). The KRAS G-domain (residues 1–166), including the conserved nucleotide-binding pocket and Switch Ⅰ/Ⅱ regions, was clearly resolved. Both the *F*_o_–*F*_c_ map and the anomalous difference Fourier map identified a single Au atom associated with each KRAS G12C monomer, whereas no corresponding anomalous density was observed in the native KRAS G12C structure solved at 2.0 Å (**Figure 5a, Figure S13a**). The Au atom was localized at the mutant Cys12 residue, with an Au–S distance of 2.3 Å, consistent with direct Au–S coordination, and the local coordination environment also involved Gly13 (**Figure 5b, Figure S13b**). Additional interactions involving GDP and bound water molecules were observed within this coordination environment (**Table S7**), supporting stabilization of the GDP-bound KRAS complex.

**Figure 5.**
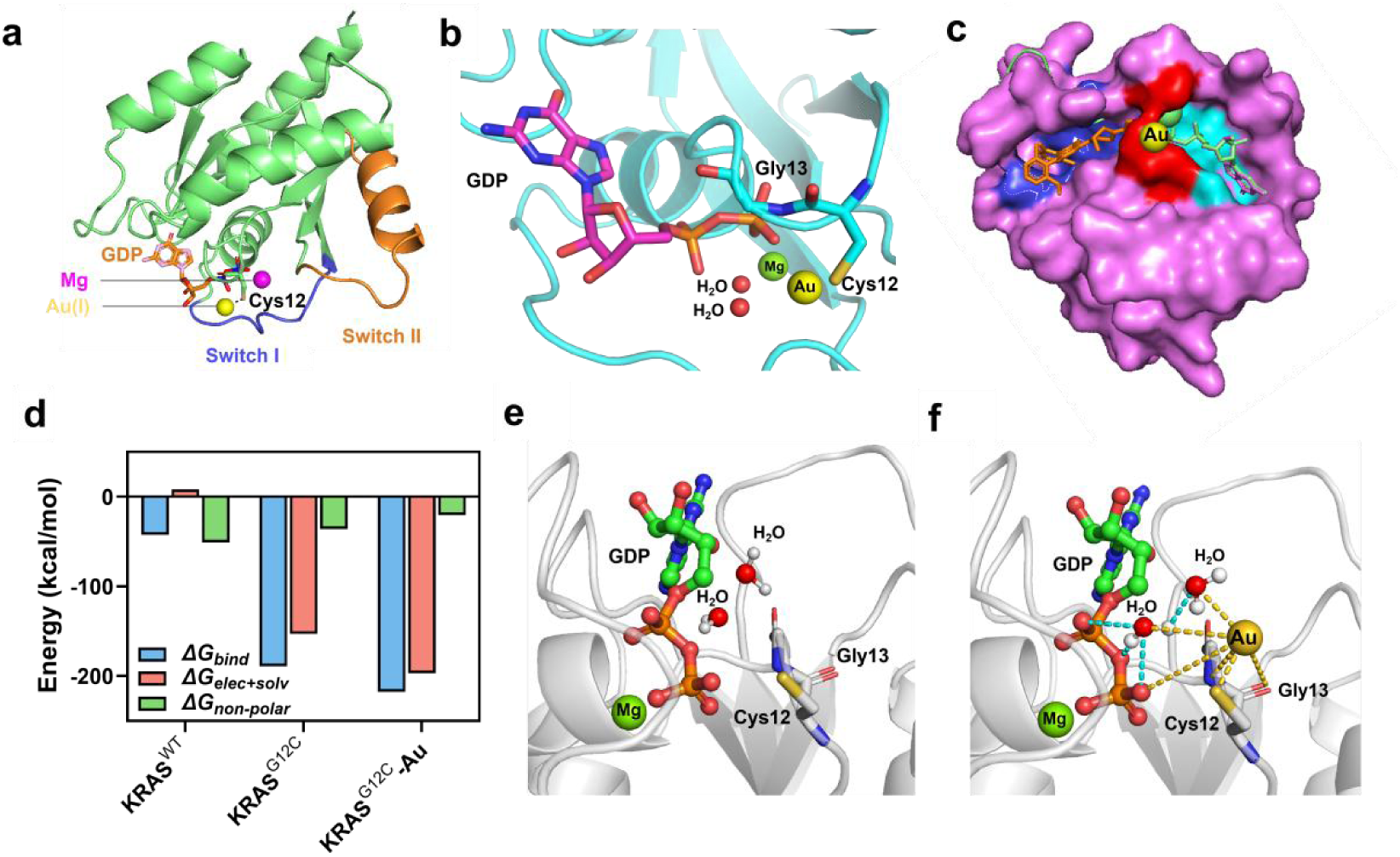
Single-atom Au engages Cys12 of KRAS G12C and stabilizes an inactive state. **(a)** Structure of Au-bound KRAS G12C showing the localization of Au at Cys12. Cys12 is shown as sticks, and the Au–S interaction is indicated by a dashed line (distance, 2.3 Å). **(b)** Enlarged view of the Au coordination environment, showing Cys12, Gly13 and GDP as sticks, Au as a yellow sphere and Mg as a green sphere. **(c)** Superposition of the Au-bound KRAS G12C structure with sotorasib (AMG510)-bound KRAS G12C (PDB 6OIM). Sotorasib is shown as orange sticks and occupies the inducible Switch-II pocket (blue), whereas Au is localized within a spatially confined coordination environment (red) around Cys12 adjacent to the nucleotide-binding pocket (cyan). **(d)** Predicted interaction energies between GDP and wild-type KRAS, KRAS G12C or Au-bound KRAS G12C. **(e, f)** Predicted binding modes of GDP with KRAS G12C (e) and Au-bound KRAS G12C (f) from MD simulations. Predicted Au-associated interactions are indicated by yellow dashed lines and water-mediated interactions by cyan dashed lines; these interactions are detailed in **Table S7**.

Comparison of the Au-bound structure with GDP-bound KRAS G12C (PDB: 4LDJ) revealed conformational differences predominantly in the Switch I region (**Figure S13c**). By contrast, the covalent KRAS G12C inhibitor sotorasib (AMG510; PDB: 6OIM) binds within the inducible Switch-II pocket (SIIP) adjacent to Cys12 (**Figure 5c**). Whereas clinically developed KRAS G12C inhibitors[22], including sotorasib[23–25], adagrasib[26, 27], glecirasib[28], and the most recent divarasib[29, 30] exploit this extended ligand-binding pocket, the Au atom is localized within a spatially confined coordination environment around Cys12 at the edge of the nucleotide-binding pocket (**Figure 5c, Figure S13d**). Therefore, as observed for EGFR, single-atom Au engages KRAS through a highly localized coordination environment rather than occupying an extended small-molecule binding pocket.

MD simulations further suggested that Au binding strengthened the predicted non-bonded interaction between GDP and KRAS G12C (**Figure 5d**). This difference was associated with additional interactions introduced by Au coordination, together with water-mediated hydrogen bonds involving GDP (**Figure 5e,f**), supporting stabilization of the GDP-bound complex and potentially restricting GDP–GTP exchange. Collectively, the EGFR and KRAS structural analyses support a common mechanism in which single-atom Au engages critical cysteine residues within spatially confined coordination environments adjacent to nucleotide-binding pockets, thereby stabilizing nucleotide-bound inactive states.

### 10714–Au suppresses tumor growth in NSCLC xenograft and patient-derived organoid models

Having established the targeting, cellular activity and molecular mechanism of 10714–Au, we finally tested whether these properties translate into antitumor efficacy *in vivo* and in patient-derived NSCLC models. Pharmacokinetic analysis in rats showed sustained plasma exposure to both 10714–Au and GE11–Au, with half-lives of approximately 15–24 h (**Figure S14**). We then assessed the antitumor effects of 10714–Au and GE11–Au at 6 or 10 mg kg⁻¹ in H1975 (EGFR L858R/T790M) and H2030 (KRAS G12C) xenograft models. Both molecules inhibited tumor growth in a dose-dependent manner, with 10714–Au at 10 mg kg⁻¹ showing the greatest effect; this was also reflected in final tumor size and weight (**Figure 6a–d**). H&E staining of excised tumors showed reduced tumor cellularity, while TUNEL staining demonstrated increased tumor cell death (**Figure S15**). Consistent with the cellular signaling analyses, immunofluorescence staining showed dose-dependent reductions in phosphorylated EGFR, AKT and ERK in H1975 tumors and in phosphorylated AKT and ERK in H2030 tumors, with the greatest suppression following treatment with 10714–Au at 10 mg kg⁻¹ (**Figure S16**). ICP-MS analysis further showed greater intratumoral Au accumulation following treatment with 10714–Au than with GE11– Au (**Figure 6e,f**). Both molecules also showed substantial renal Au accumulation, indicating shared primary excretion via glomerular filtration. Treatment was well tolerated under the conditions examined, with no significant changes in body weight or evident histopathological abnormalities in major organs (**Figure S17, S18**), while hematological and serum biochemical parameters remained within normal ranges (**Table S8, S9**).

**Figure 6.**
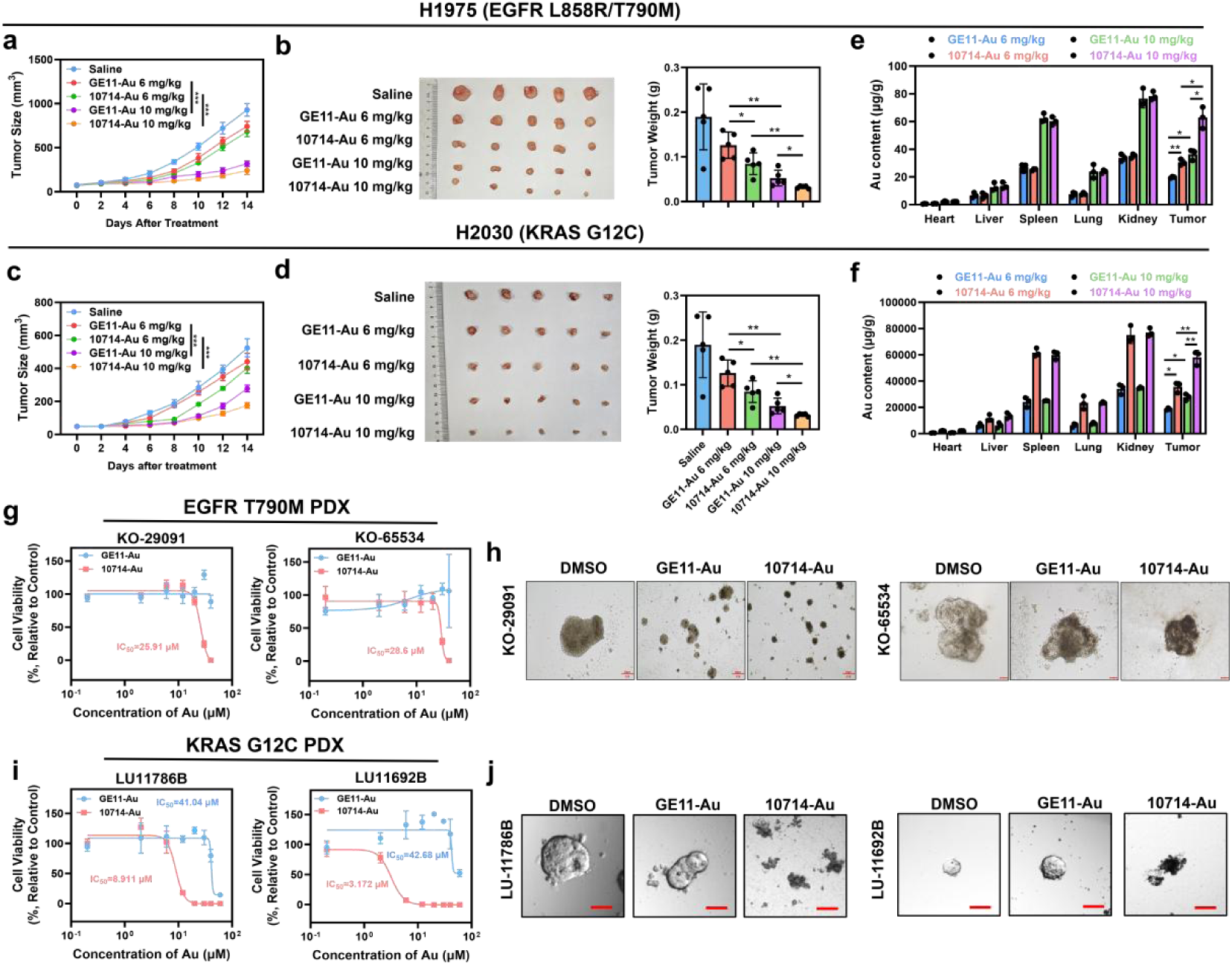
10714–Au suppresses tumor growth in NSCLC xenograft and patient-derived organoid models. **(a)** Tumor growth curves for H1975 xenografts following the indicated treatments. **(b)** Final tumor weights and representative images of excised H1975 tumors. **(c)** Tumor growth curves for H2030 xenografts following the indicated treatments. **(d)** Final tumor weights and representative images of excised H2030 tumors. **(e,f)** Au biodistribution in major organs and tumors from H1975 (e) and H2030 (f) xenograft models, measured by ICP-MS. **(g)** Viability of two EGFR T790M PDOs (KO-29091 and KO-65534) following treatment with the indicated concentrations of 10714–Au or GE11–Au. **(h)** Representative images of EGFR T790M PDOs following the indicated treatments. Scale bar, 100 μm. **(i)** Viability of two KRAS G12C PDOs (LU11786B and LU-116932B) following treatment with the indicated concentrations of 10714–Au or GE11–Au. **(j)** Representative images of KRAS G12C PDOs following the indicated treatments. Scale bar, 100 μm.

To extend these findings to patient-derived models, we tested both compounds in five NSCLC patient-derived organoids (PDOs), including two harboring EGFR T790M and three harboring KRAS G12C (**Table S10**). Across these models, 10714– Au consistently showed greater inhibitory activity than GE11–Au, extending its superior activity in cell-line and xenograft models to NSCLC PDOs (**Figure 6g–j, Figure S19**). Together, these findings demonstrate that the molecular and cellular activity of 10714–Au translates into antitumor efficacy across genetically defined *in vivo* models and NSCLC PDOs, supporting the translational potential of the targeted single-atom Au strategy.

## Discussion

Conventional inhibitor design relies on molecular recognition within suitable binding pockets, creating a long-standing challenge for targeting oncogenic proteins that lack readily druggable structural environments. Here, we establish single-atom Au as an inhibitor of two structurally distinct oncogenic drivers by exploiting the intrinsic chemical reactivity of Au towards cysteine residues. In EGFR T790M and KRAS G12C, single-atom Au engaged Cys797 and Cys12, respectively, within spatially confined coordination environments adjacent to the nucleotide-binding pockets, with structural and computational analyses supporting stabilization of inactive nucleotide-bound states. Thus, this cysteine-targeting mechanism provides a metal-based strategy for oncogenic-protein inhibition that relies on highly localized atomic coordination rather than engagement of an extended ligand-binding pocket.

The development of a peptide-based delivery system was critical for translating this chemical concept into *in vivo* efficacy. Conventional approaches to peptide discovery typically rely on screening large experimental libraries, for example by phage display or chemical library screening, or increasingly on large-scale computational approaches. However, the limited number of experimentally validated EGFR-targeting peptides constrains conventional data-intensive computational strategies. Our few-shot learning approach, built on a pretrained LSTM architecture and fine-tuned using only 54 experimentally validated EGFR-targeting peptides, integrated machine learning-based sequence generation with structure-based physical filtering to identify novel candidate peptides with high predicted EGFR-binding affinity. Importantly, peptide 10714 showed greater EGFR-binding affinity than the experimentally validated GE11 comparator, while 10714–Au showed greater cellular Au uptake than GE11–Au and greater activity across the biochemical, cellular and *in vivo* models examined. This approach therefore exemplifies how few-shot learning can accelerate the discovery of functional targeting peptides when experimental training data are scarce and could potentially be extended to other receptor-targeted delivery systems.

The structural insights from our crystallographic studies reveal a striking distinction between single-atom Au and conventional covalent inhibitors. In EGFR, osimertinib occupies an extended region of the ATP-binding pocket and engages Cys797 through its acrylamide warhead[31]. KRAS, by contrast, was historically considered difficult to target, until the identification of the inducible Switch-II pocket (SIIP) enabled the development of covalent KRAS G12C inhibitors, including sotorasib and adagrasib, which exploit the GDP-bound state to position an electrophilic warhead for engagement of Cys12[24, 26]. In both cases, these small-molecule inhibitors use an extended molecular scaffold to establish interactions within a defined binding pocket and appropriately position a reactive group for cysteine engagement. By contrast, a single Au atom can exploit its intrinsic preference for thiolate coordination within a much more spatially confined environment. Consistent with this principle, Au was localized adjacent to the respective nucleotide-binding pockets of both EGFR T790M and KRAS G12C, coordinated by Cys797 and Cys12, respectively. Together with the additional interactions identified in the structural and computational analyses, these localized coordination environments support stabilization of nucleotide-bound inactive states. These findings therefore suggest a distinct inhibitor-design principle in which target engagement can be achieved through highly localized atomic coordination rather than requiring the extended molecular recognition interface of a conventional small-molecule ligand.

An important challenge for targeted cancer therapy is the emergence or coexistence of multiple oncogenic alterations, which can necessitate targeting more than one driver[1, 32]. The structural divergence of target proteins can make such multi-target inhibition difficult to achieve within a single conventional small-molecule scaffold[33]. Combined or sequential use of different targeted inhibitors can also introduce additional toxicity, cost and pharmacokinetic complexity[34]. The ability of single-atom Au to engage cysteine residues within highly localized coordination environments in two structurally distinct oncogenic proteins therefore raises the possibility that this strategy could ultimately enable multi-target inhibition without requiring a separate molecular recognition scaffold for each target.

Several limitations and open questions warrant further investigation. First, although we have demonstrated activity against EGFR T790M and KRAS G12C, the molecular selectivity and generality of this mechanism remain to be established. Given the intrinsic affinity of Au for thiol groups and the widespread occurrence of cysteine residues across the proteome, it will be important to define the broader intracellular target profile of Au and determine the structural and chemical features that distinguish functionally susceptible cysteine environments from other potential Au-binding sites. Such studies will also help establish whether single-atom Au inhibition can be selectively extended to other cysteine-containing oncogenic proteins. Second, EGFR and KRAS mutations rarely co-occur in the same patient, and our current preclinical models predominantly represent tumors driven by individual oncogenic mutations[35]. Whether the single-atom Au strategy can simultaneously inhibit multiple targets in tumors harboring concurrent or sequential oncogenic alterations therefore requires direct investigation. Finally, future studies should establish the pharmacological properties, long-term safety and therapeutic window of 10714–Au and determine whether single-atom Au inhibition can be effectively combined with existing treatments, including chemotherapy or immunotherapy.

In conclusion, this work establishes a strategy for single-atom inhibition of oncogenic drivers through cysteine-targeting Au coordination chemistry. The combination of AI-driven peptide discovery and single-atom coordination provides a framework that could be extended to other cysteine-bearing oncogenic proteins as the determinants of Au susceptibility and selectivity are defined. More broadly, the single-atom inhibitor concept may expand how targetability is considered in drug discovery: beyond the geometry of conventional ligand-binding pockets to include appropriately positioned cysteine residues and local environments capable of supporting atomic coordination.

## Materials and Methods

### Few-shot learning-based generation of EGFR-targeting peptides

A LSTM-based peptide-generation model was pretrained to learn general amino acid patterns and sequence dependencies and subsequently fine-tuned using a dataset of 54 experimentally validated EGFR-targeting peptides[36]. Candidate sequences were generated through iterative mutation and resampling while preserving sequence features learned from the validated peptides.

Approximately 12,000 candidate peptides were generated in each independent generation batch. The generated and experimentally validated peptides were projected into the same chemical-feature space using PCA. Candidate peptides located within or near the chemical space occupied by the experimentally validated peptides were retained for subsequent structural prediction, molecular docking, and MD screening. Peptides were next screened using molecular docking and MD simulations. The crystal structure of EGFR was obtained from the Protein Data Bank (PDB; PDB ID: 1NQL). ZDOCK was first used to explore possible binding modes across translational and rotational space between EGFR and the peptides and to rank the resulting docking poses. A total of 753 generated peptides were selected for further screening. AutoDock CrankPep (ADCP) was subsequently used for blind docking of the selected peptides to EGFR. Each ADCP calculation comprised 50 independent searches of 2.5 million Monte Carlo steps and generated the 10 highest-ranked binding modes. The top 20 peptides were selected for MD simulations.

MD simulations were performed using GROMACS. The AMBER ff14SB force field was used to describe the protein and peptides, and the TIP3P water model was used for solvation. MD trajectories were analysed using Visual Molecular Dynamics (VMD), which was also used to calculate non-bonded interaction energies between EGFR and the peptides. Effective binding energies of the protein–peptide complexes were calculated using gmx_MMPBSA. More negative values indicate more favourable predicted interactions. Five peptides were selected for subsequent experimental evaluation, since they ranked top 5 by non-bonded interaction energies.

### Preparation and characterization of peptide-Au molecules

Using GE11–Au as an example, 5 mg of GE11 (YHWYGYTPQNVIKKKKYCC) was dissolved in 2 mL of ultrapure water. NaOH solution (0.5 M, 100 μL) was then added to adjust the pH 12.0. The reaction was conducted at 42°C. After stirring for 3 min, HAuCl_4_ (25 mM, 40 μL) was added with vigorous stirring, and the mixture was incubated for 12 h. The reaction mixture was subsequently stored in the dark at 4°C for 2 days. Before use, the product was dialysed using a 3-kDa molecular-weight-cutoff (MWCO) dialysis membrane (Millipore) to remove free peptide and other ions. The purified product was stored at 4°C in the dark until further use. The other five peptide–Au molecules were prepared using identical reaction conditions and stoichiometric ratios.

Fluorescence spectra were acquired using a spectrofluorometer (RF-5301, Shimadzu, Japan). Molecular composition was analysed by MALDI-TOF-MS using an SA matrix in linear positive-ion mode. UV–visible (UV–Vis) absorption spectra were acquired using a UV-1800 spectrophotometer (Shimadzu, Japan). Particle size distributions and ζ-potentials were measured using a Zetasizer Nano ZS90 (Malvern, UK). The concentrations of the peptide–Au molecules were determined by an ICP-MS analysis system (Thermo Elemental X7, USA).

### Conformational search and structural optimization of the Au_5_Peptide_3_ complex

An initial molecular model of the Au₅Peptide₃ complex was constructed according to the Au-to-peptide stoichiometry determined by MALDI-TOF-MS. The sulfur atoms of the YCC motifs were assigned as coordination sites. Global conformational sampling was performed using ABCluster[37]. A total of 100 candidate conformations of the Au₅Peptide₃ complex were generated and subsequently subjected to full geometry optimization using the GFN1-xTB semiempirical tight-binding method.^40^ The electronic energies of the optimized structures were compared, and the structure with the lowest GFN1-xTB energy was selected as the representative configuration for structural analysis and visualization.

### Surface plasmon resonance (SPR) for affinity studies

SPR measurements were performed using a Biacore 8K+ instrument (GE Healthcare, Piscataway, NJ, USA) at 25°C. The proteins were immobilized on a CM5 sensor chip by amine coupling. PBST (PBS containing Tween-20) was used as the running buffer.

For EGFR-targeting efficiency evaluation (**Figure 2g, h, Figure S4**), purified extracellular EGFR (ab155639, Abcam) was employed. Peptides and peptide–Au molecules were diluted to the indicated concentrations in PBST and injected over the immobilized proteins to obtain SPR sensorgrams. For peptides, contact time was 60s, while dissociation time was 60s. The flow rate was 30 μL/min. For peptide-Au molecules, contact time was 120s, while dissociation time was 360s. The flow rate was 30 μL/min.

For the binding affinity assessment to the intracellular domain (containing the enzymatic domain) of EGFR, intracellular wild-type EGFR (HY-P72987, MCE) and intracellular EGFR L858R/T790M (HY-P702832, MCE) were adopted (**Figure S6**). Peptide–Au molecules were diluted to the indicated concentrations in PBST and injected over the immobilized proteins to obtain SPR sensorgrams. Contact time was 90s, while dissociation time was 180s. The flow rate was 30 μL/min.

For measuring the binding to KRAS G12C, recombinant human KRAS G12C (ab314426, Abcam) was used (**Figure S7**). Peptide–Au molecules were diluted to the indicated concentrations in PBST and injected over the immobilized proteins to obtain SPR sensorgrams. Contact time was 90s, while dissociation time was 180s. The flow rate was 30 μL/min.

Reference-subtracted sensorgrams were analyzed using Biacore Insight Evaluation software version 3.0. Based on the rapid association and dissociation characteristics of the sensorgrams, the data were fitted using a steady-state affinity model to determine the equilibrium dissociation constant (*K*_D_).

For regeneration conditions of all SPR experiments, glycine (pH 2.0) was used, contact time was 30s, and the flow rate was 30 μL/min.

### Oncogenic driver activity assays

EGFR kinase activity for wild-type, L858R/T790M, or C797S EGFR was measured mainly using the EGFR (T790M/L858R) Kinase Assay (V5325, Promega, USA) according to the manufacturer’s instructions, while EGFR wild-type or EGFR C797S (ab208478, Abcam) proteins were purchased separately. Different concentrations of 10714–Au or GE11–Au were incubated with intracellular wild-type, L858R/T790M, or C797S EGFR (ab208478, Abcam).

KRAS nucleotide exchange activity was measured using the RAS GEF Exchange Assay (BK101, Cytoskeleton) according to the manufacturer’s instructions. Different concentrations of 10714–Au or GE11–Au were incubated with wild-type KRAS (ab268714, Abcam) or KRAS G12C.

### Cells

H1975, H292, H522, H1299, H2122, H358, and H2030 cells were purchased from Procell (Wuhan, China) and cultured in high-glucose Dulbecco’s modified Eagle medium (DMEM) supplemented with 10% heat-inactivated fetal calf serum and 1% (v/v) penicillin–streptomycin at 37℃ in a humidified atmosphere containing 5% CO_2_. Cells were routinely tested for mycoplasma contamination. Cells were used for experiments during logarithmic growth.

### Cellular uptake route of 10714-Au

For confocal microscopy, H1975 cells were seeded onto coverslips in 24-well plates and incubated overnight. Cells were pretreated with DMSO, amiloride (100 μM), Pitstop 2 (10 μM), methyl-β-cyclodextrin (MβCD; 1 mg/mL), or nystatin (20 μg/mL) for and subsequently incubated with 10714–Au (60 μM) for 24 h. Cells were then washed three times with PBS and examined by confocal microscopy.

For ICP-MS analysis, cells (1 × 10^5^ cells/well) were seeded in 6-well plates and treated under the same conditions. Cell numbers were determined before the samples were digested overnight in aqua regia. The aqua regia was subsequently evaporated by heating, and the samples were reconstituted in an aqueous solution containing 2% nitric acid and 1% hydrochloric acid. Au content was measured by ICP-MS, and divided by cell number to obtain Au-per-cell values.

### Lysosomal escape of 10714-Au

H1975 cells were seeded in glass-bottom culture dishes and cultured for 12 h. Cells were then incubated with 10714–Au (60 μM) for 0, 0.5, 6, or 24 h. Lysosomes were stained with LysoTracker Red (HY-D1300, MCE) for 20 min, after which the cells were washed and examined by confocal microscopy to assess colocalization of 10714–Au with lysosomes. Pearson’s correlation coefficient (r) was calculated using ImageJ.

For ICP-MS analysis, H1975 cells were cultured in T75 flasks and treated with 10714–Au (60 μM) for 0, 0.5, 6, or 24 h. Cell numbers were determined, and lysosomes were subsequently isolated using a lysosome extraction kit (bb-3603, BestBio, China). Samples were processed for ICP-MS analysis as described above, and Au content per cell of lysosomal and non-lysosomal fractions was measured by ICP-MS normalized to cell number.

### Western blotting

Cells were lysed in RIPA lysis buffer containing a protease and phosphatase inhibitor cocktail (P1051, Beyotime, China). Protein concentrations were determined using a BCA protein assay kit (Beyotime, China). Total protein (30 μg per sample) was separated by SDS-PAGE and transferred to PVDF membranes. Membranes were blocked for 1 h at room temperature and incubated overnight with diluted primary antibodies (1:1000) against EGFR (ab52894, Abcam, UK), p-EGFR (ab40815, Abcam, UK), AKT (ab300473, Abcam, UK), p-AKT (29163, Proteintech, USA), ERK (ab184699, Abcam, UK), p-ERK (ab214036, Abcam, UK), and GAPDH (ab8245, Abcam, UK) at 4°C. Membranes were then incubated with the appropriate secondary antibodies for 1 h at room temperature, and immunoreactive proteins were detected using Amersham ECL Prime Western Blotting Detection Reagent.

### Cell viability and apoptosis assays

Cell viability was assessed using the Cell Counting Kit-8 (CCK-8) assay. Cells (1 × 10^5^ cells/mL) were seeded in 96-well plates at 100 µL per well and treated with peptide–Au molecules or the corresponding peptides at a range of concentrations for 24 h. Cell viability was then measured using CCK-8 reagent (Beyotime, China).

Apoptosis was assessed through Annexin V–FITC and propidium iodide (PI) staining and flow cytometry. H1975 or H292 cells (1 × 10^5^ cells/well) were treated with a range of concentrations of 10714–Au or GE11–Au for 24 h. Cells were resuspended in 490 μL of Annexin V–FITC binding buffer, followed by the addition of 5 μL of Annexin V–FITC and incubation for 10 min (C1062L, Beyotime). PI solution (10 μL) (HY-D0815, MCE) was then added, and the cells were incubated in the dark for 20 min. Samples were analyzed using an Accuri C6 flow cytometer.

### X-ray crystallography

The T790M/V948R-mutant EGFR kinase domain (residues 696–1022) was expressed using a baculovirus/Sf9 insect-cell expression system. The lysis buffer contains 20 mM Tris-HCl pH 8.0, 150 mM NaCl, 3 mM KCl, 1% glycerol, 1 mM PMSF, 1 mM TCEP, 20 mM imidazole added with a protease inhibitors cocktail. The protein was purified by Ni^2+^ affinity chromatography followed by size-exclusion chromatography using a Superdex 200 column with running buffer containing 20 mM Tris (pH 8.0), 150 mM NaCl, 1% glycerol, and 0.5 mM TCEP.

KRAS G12C was cloned into a pET28(+) expression vector and expressed in Escherichia coli BL21(DE3) cells. The construct contained an N-terminal His_6_-tag followed by a TEV protease cleavage site. Protein expression was induced with 0.2 mM isopropyl β-D-1-thiogalactopyranoside (IPTG) at 16°C for 14 h. Cells were then resuspended in lysis buffer (20mM Tris/HCl, pH 8.0, 20 mM imidazole and 300 mM NaCl) and lysed by French press, followed by centrifuge at 15000 r for 50 min. The supernatant was loaded onto a Ni-NTA column pre-equilibrated with lysis buffer and eluted with 20 mM Tris-HCl (pH 8.0), 300 mM NaCl, and 300 mM imidazole. The buffer was subsequently exchanged to 20 mM Tris (pH 8.0), 150 mM NaCl, 5 mM imidazole, 5 mM MgCl_2_, and 1 mM β-mercaptoethanol, supplemented with 1 mg GDP per 20 mg protein. The His6-tag was cleaved with TEV protease at 4°C, after which the TEV protease was removed by reverse Ni-NTA affinity chromatography. The protein was then further purified by size-exclusion chromatography as described for EGFR.

EGFR crystals were grown at 20°C by sitting-drop vapor diffusion using 0.7 μL of EGFR protein solution (10 mg/mL) mixed with an equal volume of reservoir solution containing 0.1 M Bis-Tris (pH 5.0), 22.5% PEG 3350, and 5 mM TCEP, in the presence of 1 mM AMP-PNP (A) and 10 mM MgCl₂. AMP-PNP was used to substitute ATP because structures of EGFR in complex with ATP are rarely obtained. KRAS G12C crystals were grown at 20°C by sitting-drop vapor diffusion using 0.6 μL of protein solution (25 mg/mL) mixed with 0.7 μL of reservoir solution containing 0.2 M ammonium chloride and 22% PEG 3350. EGFR and KRAS crystals were soaked for >24 h in their respective reservoir solutions supplemented with 10 mM 10714–Au, cryoprotected, and flash-frozen in liquid nitrogen.

Diffraction data for the EGFR–Au complex were collected at 100 K at beamline BL02U1 of the Shanghai Synchrotron Radiation Facility (SSRF), and diffraction data for the KRAS G12C–Au complex were collected at beamline BL10U2. Diffraction data were processed using XDS. Initial phases were determined by molecular replacement using Phaser in the CCP4 suite, with metal-free EGFR T790M/V948R or KRAS G12C structures as the initial models. Model building was performed using Coot, and structure refinement was performed using Phenix and Refmac. MolProbity was used for structure validation. The location of the Au ion was identified from the *Fo* − *Fc* and anomalous difference Fourier maps. Data collection, refinement, and validation statistics are provided in Table S5. Structural figures were prepared using PyMOL (http://www.pymol.org).

### Molecular dynamics simulations

Experimentally determined Au-bound EGFR T790M/V948R and Au-bound KRAS G12C structures were used as starting structures for MD simulations. Au-free control systems were generated by removing the Au atom while retaining the same initial protein coordinates, nucleotides, Mg²⁺ ions, and structurally relevant crystallographic water molecules. The EGFR crystal structure contained AMP-PNP. For the ADP-bound MD system, ADP was generated by removing the γ -phosphate group from AMP-PNP while retaining the coordinates of the remaining nucleotide atoms. The crystallographic GDP molecule was retained in the KRAS G12C simulations. Au-free control systems were generated by removing Au while retaining the remaining atomic coordinates. Proteins were described using the AMBER ff14SB force field,[38] whereas nucleotides and other nonstandard components were parameterized using GAFF, Antechamber, and AM1-BCC charges.[39, 40]

The Au–S coordination between Au and Cys797 in EGFR or Cys12 in KRAS was explicitly represented using bonded force-field terms in the MCPB.py model.^45^ In contrast, Au···O contacts and other noncovalent interactions were not introduced as bonded terms and were identified from subsequent trajectory analyses. A reduced metal-center model containing Au, the coordinating cysteine residue, and the surrounding coordination environment was optimized using ORCA version 5.0.4 at the TPSS/def2-TZVP level.[41–44] For Au, the effective core potential associated with the def2 basis-set family was applied to describe the inner-shell electrons. The optimized geometries and calculated Hessian matrices were used to derive equilibrium Au–S bond lengths, Au-centered equilibrium angles, and the corresponding bonded force constants.

Each system was solvated in TIP3P water and neutralized with counterions. Following energy minimization, the systems were equilibrated for 5 ns under NVT conditions and then for 5 ns under NPT conditions at 310.15 K and 1 bar. Unrestrained 50-ns production simulations were performed using GROMACS with a 2-fs time step.[45] One independent 50 ns production trajectory was generated for each system. Trajectory stability and interactions were evaluated using backbone root-mean-square deviation (RMSD), residue-level root-mean-square fluctuation (RMSF), nucleotide–pocket interactions, Au–S distances, Au···O contacts defined using an Au– O distance cutoff of 5.0 Å, and hydrogen-bond occupancies identified using a donor– acceptor distance cutoff of 3.0 Å. Interaction occupancies were calculated as the percentage of analyzed trajectory frames satisfying the corresponding geometric criteria. MM/PBSA analysis was performed using gmx_MMPBSA on snapshots extracted from the 5 – 50 ns portion of each production trajectory, with every fifth trajectory frame selected for calculation. The same trajectory interval and frame-selection scheme were applied to all systems to ensure direct comparability. The nonpolar contribution (*G*nonpolar), combined electrostatic and solvation contribution (*G*elec+solv), and total effective binding energy (*G*bind) were calculated. Per-residue energy decomposition was performed to evaluate the contributions of individual residues to nucleotide binding. Conformational entropy was not included; therefore, the calculated values were interpreted as relative effective binding energies.

### Pharmacokinetic study

10714–Au or GE11-Au (5 mg/kg) was administered intraperitoneally to one male and one female Sprague–Dawley (SD) rats, respectively. Blood samples were collected from the retro-orbital venous plexus via the medial canthus at 0 h, 0.25 h, 0.5 h, 1 h, 2 h, 4 h, 8 h, 12 h, and 24 h after injection. Plasma was collected, digested, and analyzed for Au content by ICP-MS.

### Xenograft tumor model

Six-week-old female BALB/c nude mice were purchased from Beijing HFK Bioscience Co., Ltd. and maintained under standard laboratory conditions. All animal experiments were conducted in accordance with the National Law on the Use of Experimental Animals and the animal care requirements of the Ethics Committee of Beijing University of Technology (protocol no. HS202202009).

Xenograft models were established by suspending 1 × 107 H1975 cells or 1.5 × 107 H2030 cells in a 1:1 (v/v) mixture of Matrigel and PBS and injecting the cell suspension subcutaneously into the rear flank of each mouse. Once tumors reached approximately 100 mm3, mice were randomly assigned to five groups (n = 5 per group) and treated by intraperitoneal injection of 200 μL saline, GE11-Au (6 or 10 mg/kg), or 10714-Au (6 or 10 mg/kg) every 2 days. Tumor volumes was measured by by a caliper every day. The tumor volume was calculated as follows: (length × width^2^) × 0.5 and body weights were measured daily.

After 14 days of treatment, mice were euthanized, and the hearts, livers, spleens, lungs, kidneys, and tumors were collected. Tissues were either processed for ICP-MS analysis or fixed in 4% paraformaldehyde for H&E staining, TUNEL staining, and immunofluorescence analysis of the indicated target proteins. Blood samples were also collected for hematological and biochemical analyses. Whole blood was used for routine hematological analysis, and serum was used for biochemical analysis.

### Patient-derived organoids (PDOs) viability assay

Organoids were processed at a 1:1 ratio with 50% Matrigel (356231, Corning), and the mixture was mechanically sheared to generate organoids of uniform size. Organoids were then collected by adding 20 μL of 100x Dispase II solution (17105041, Gibco) and seeded into 384-well Ultra-Low Attachment microplates (4588, Corning) using a Multidrop dispenser (Thermo) at a density of 330 organoids per well. Organoids were treated with different concentrations of 10714–Au or GE11– Au for 5 days. Representative images were acquired for each treatment, and endpoint CTG measurements were obtained by Luminescent CellTiter-Glo (CTG) signals measured by the CellTiter-Glo 3D Cell Viability Assay (G9683, Promega). Briefly, 40 μL of CTG 3D reagent was added to each well, mixed for 20 min on a plate shaker, and incubated for a further 20 min at room temperature in the dark. Luminescence was then measured using an EnVision plate reader (PerkinElmer)

### Statistics

For analyses of effects of peptide–Au molecules on enzymatic activity, cell viability, and PDO growth, IC_50_ values were calculated by fitting a four-parameter variable-slope inhibitor-versus-response model in GraphPad Prism version 10.4.1. Other data are presented as the mean ± SD and were analyzed using unpaired two-tailed *t*-tests. Statistical significance was defined as *P* < 0.05. Significance is indicated as follows: \**P* < 0.05, \*\**P* < 0.01, and \*\*\**P* < 0.001; n.s., not significant.[46]

