## supplemental fig 1-19 and supplemental table 1-10 for "Single-atom inhibition of oncogenic drivers through cysteine coordination"

### Supplementary Figures

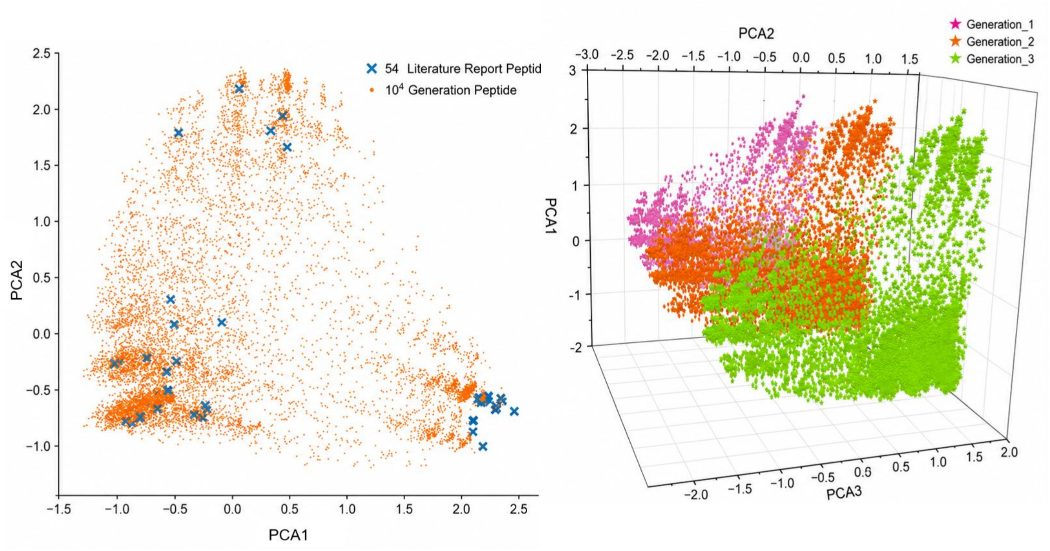


**Figure S1. PCA analysis of generated peptides. (a)** Principal component analysis (PCA) visualization of 12,000 peptides generated in a representative round of peptide generation, with the 54 literature-reported peptides indicated by blue crosses. **(b)** PCA visualization of peptides generated in three independent rounds.

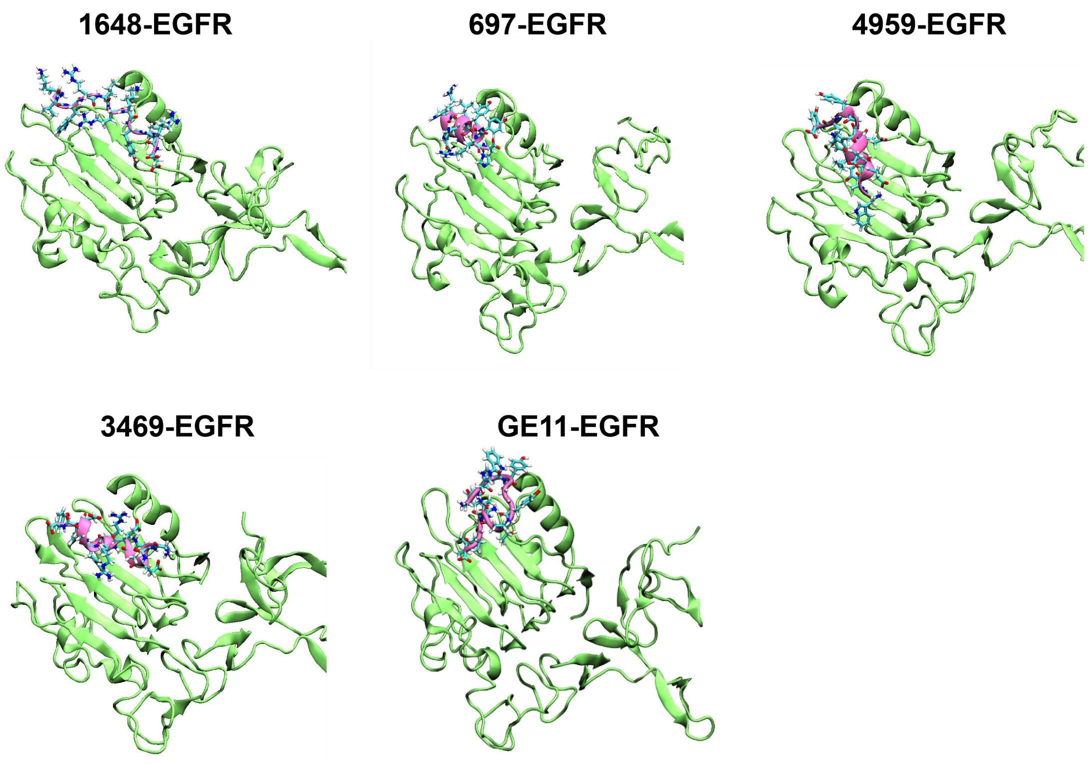


**Figure S2: MD simulation of the interactions between four additional generated peptides or GE11 and the extracellular domain of EGFR.**

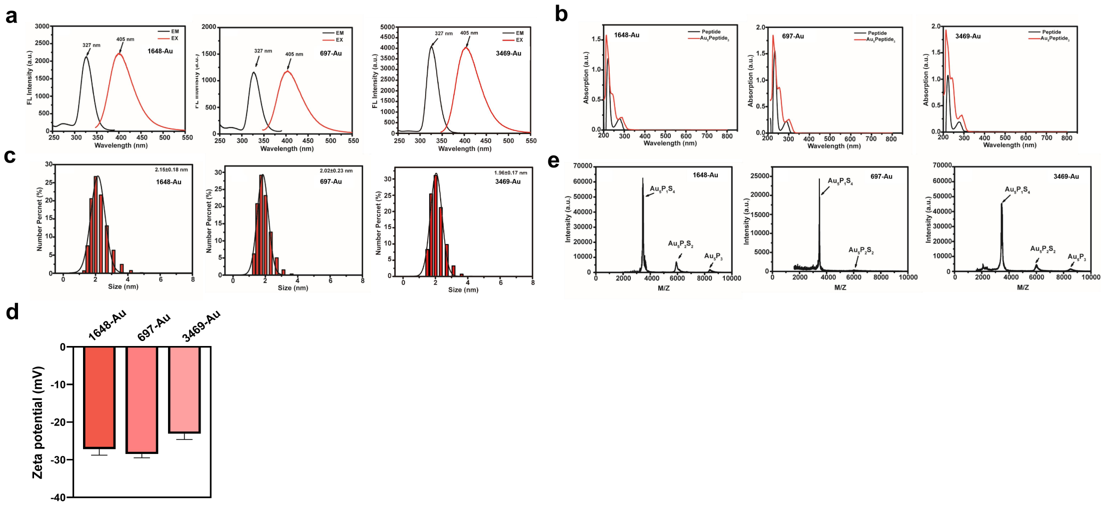


**Figure S3. Characterization of Au-delivery molecules conjugated with additional generated EGFR-targeting peptides. (a)** Fluorescence excitation (black) and emission (red) spectra. **(b)** UV–Vis spectra of the peptide–Au molecules and their respective free peptides. **(c)** Dynamic light scattering analysis. **(d)** Zeta-potential measurements. **(e)** MALDI-TOF-MS spectra acquired in positive-ion linear mode. The proposed molecular composition of each molecule is $Au_5Peptide_3$.

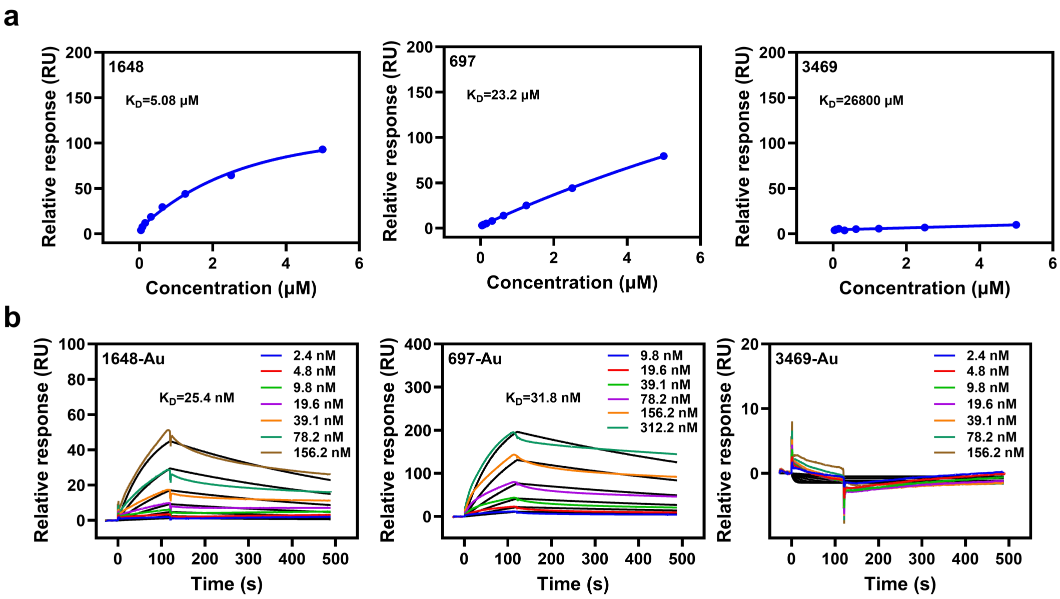


**Figure S4. SPR analysis of the binding of the indicated peptides (a) and peptide–Au molecules (b) to the extracellular domain of EGFR.**

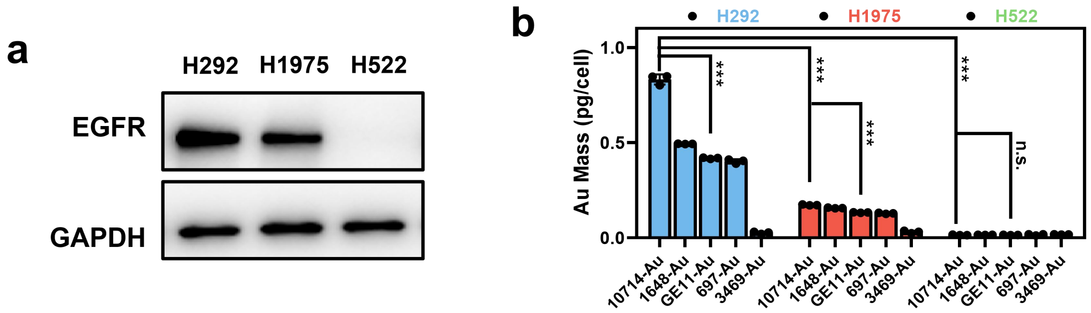


**Figure S5. Cellular Au accumulation following treatment with different peptide–Au molecules. (a)** EGFR expression in the indicated human NSCLC cell lines, assessed by western blotting. **(b)** Cellular Au accumulation following treatment with five peptide–Au molecules in three NSCLC cell lines, measured by ICP-MS.

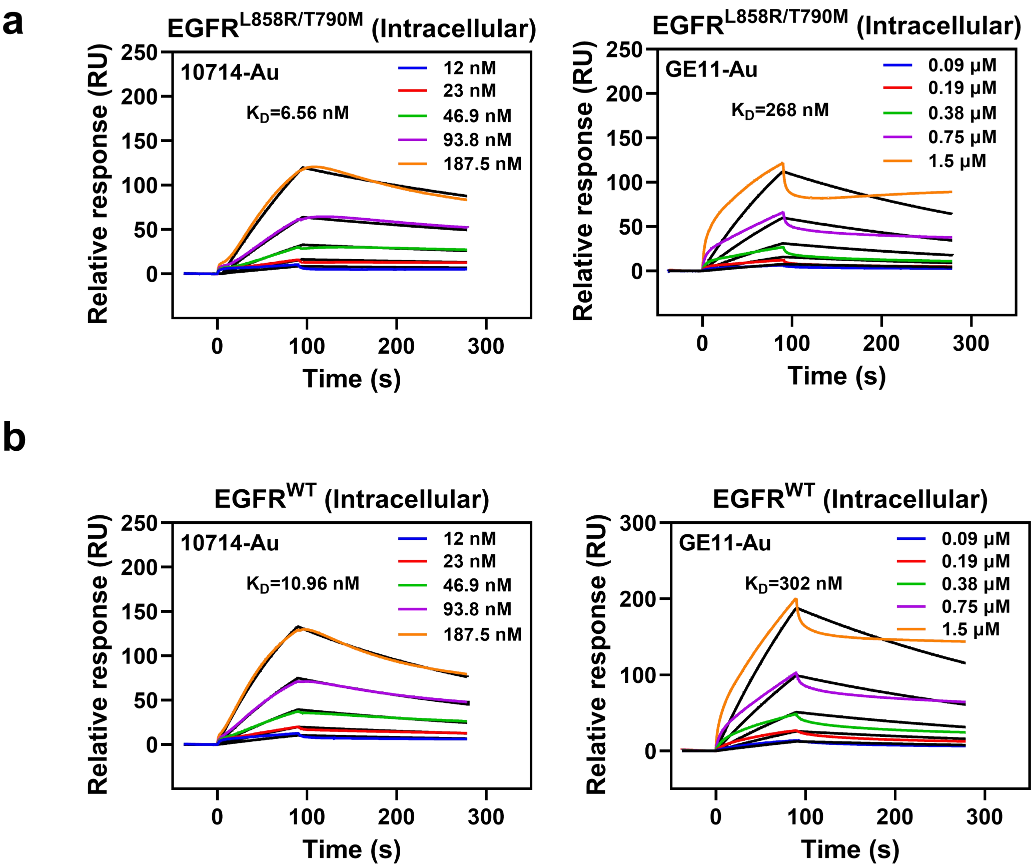


**Figure S6. SPR analysis of the binding of 10714–Au and GE11–Au to mutant (a) and wild-type (WT) (b) EGFR intracellular domains.**

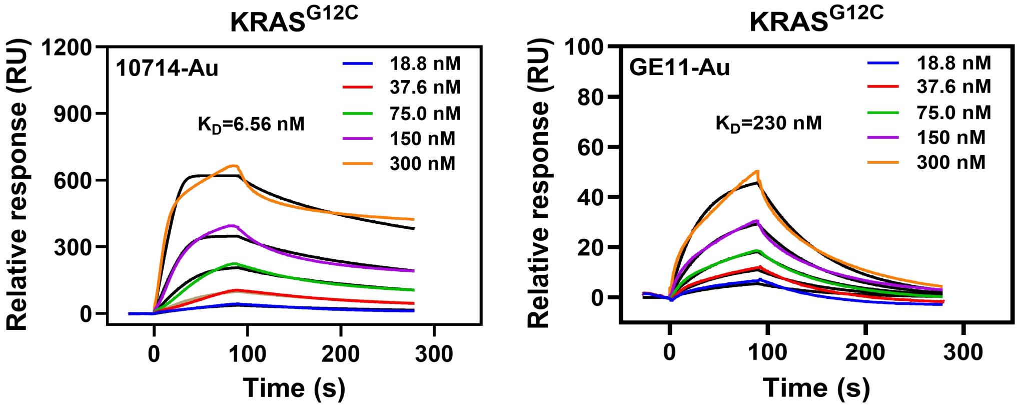


**Figure S7. SPR analysis of the binding of 10714–Au and GE11–Au to KRAS G12C.**

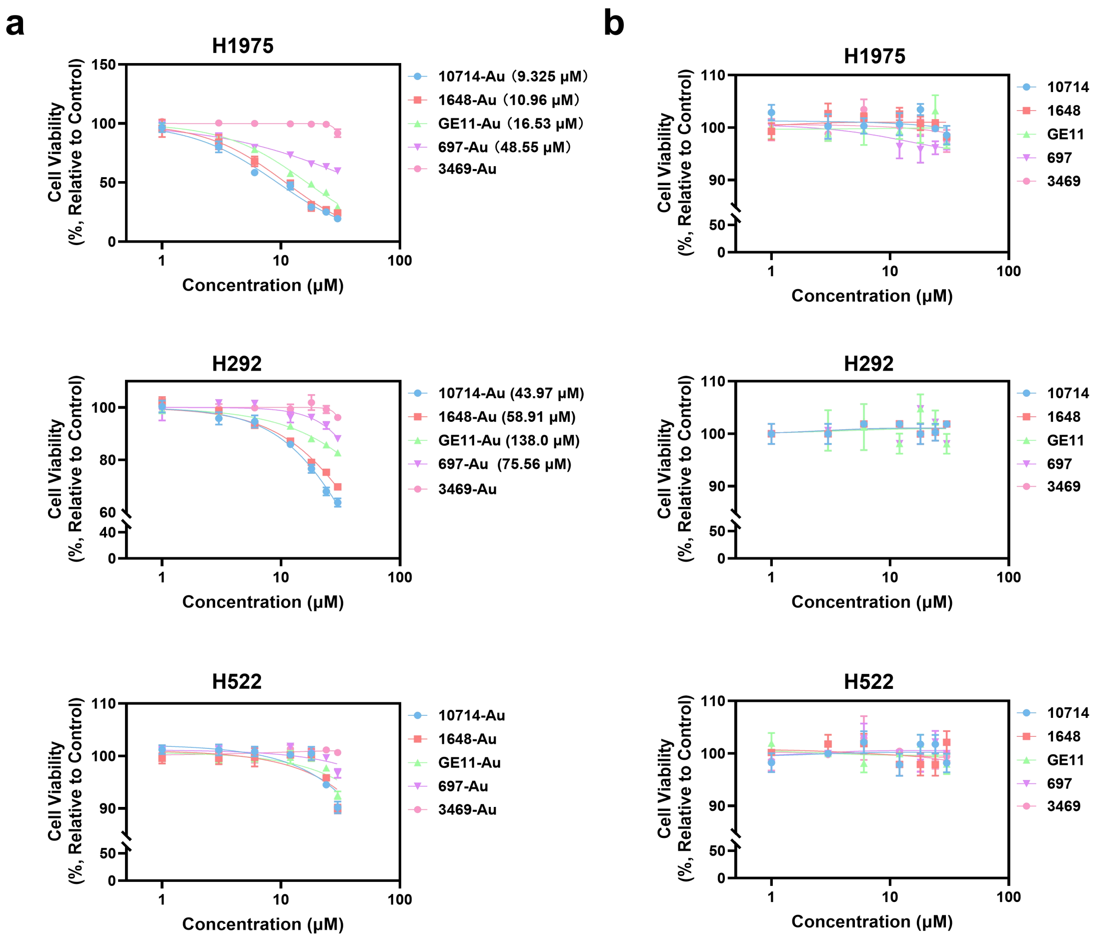


**Figure S8. Effects of peptide–Au molecules and their respective free peptides on NSCLC cell viability. (a,b)** Cell viability following treatment with the indicated peptide–Au molecules (a) or corresponding free peptides (b) in NSCLC cell lines with different levels of EGFR expression.

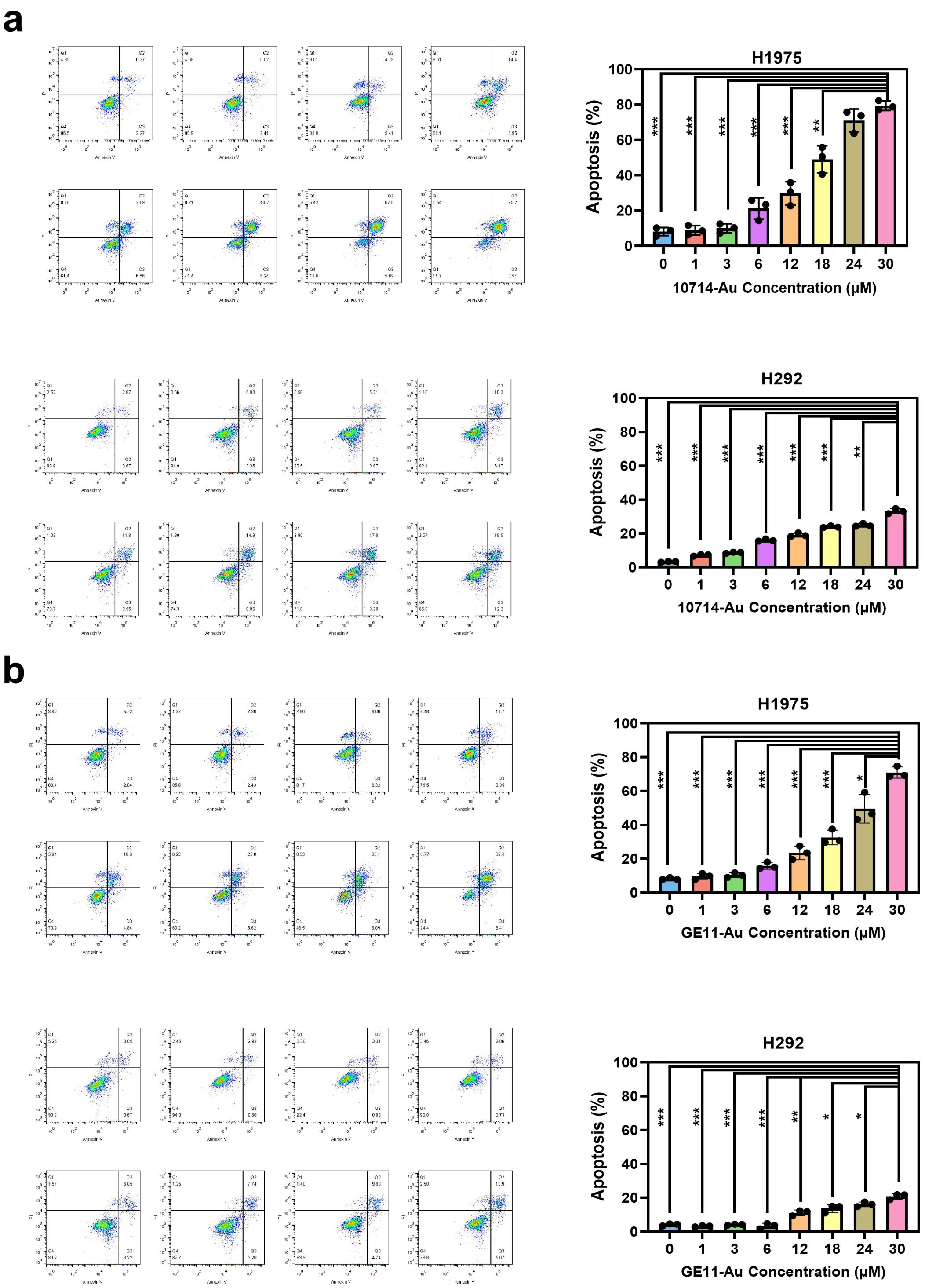


**Figure S9. Effects of 10714–Au and GE11–Au on apoptosis in NSCLC cell lines with different levels of EGFR expression.** Apoptosis following treatment with **(a)** 10714–Au or **(b)** GE11–Au was assessed by Annexin V/PI staining.

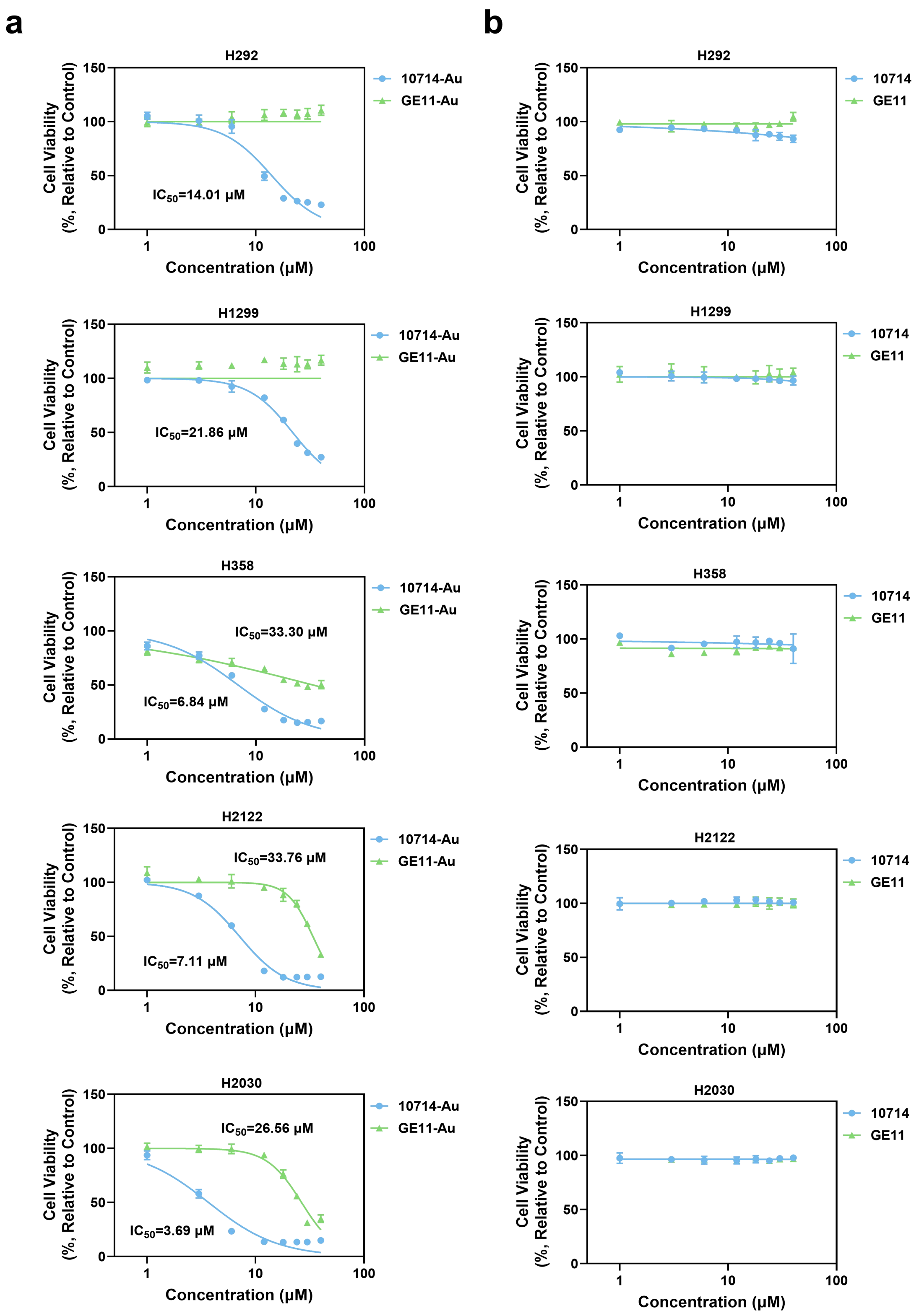


**Figure S10. Effects of peptide–Au molecules and their respective free peptides on the viability of NSCLC cells with different KRAS genotypes. (a,b)** Cell viability following treatment with the indicated peptide–Au molecules (a) or corresponding free peptides (b).

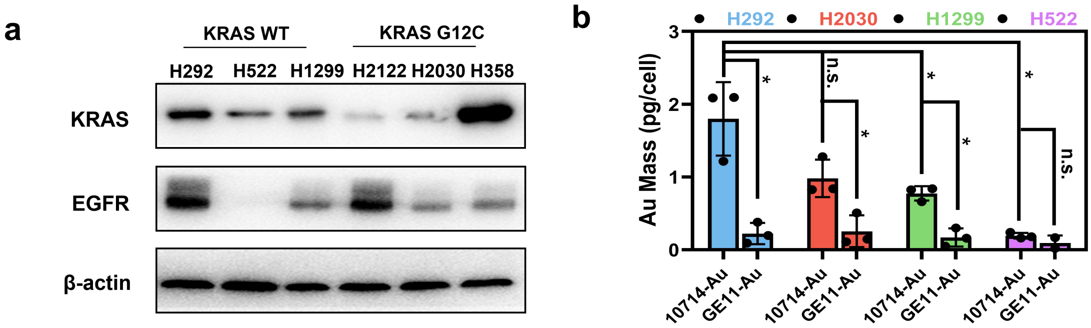


**Figure S11. EGFR expression and cellular Au accumulation in KRAS wild-type and KRAS G12C NSCLC cells. (a)** KRAS and EGFR expression in the indicated human NSCLC cell lines, assessed by western blotting. **(b)** Cellular Au accumulation following incubation with the indicated EGFR-targeting peptide–Au molecules in NSCLC cell lines with wild-type or G12C-mutant KRAS and different levels of EGFR expression, measured by ICP-MS.

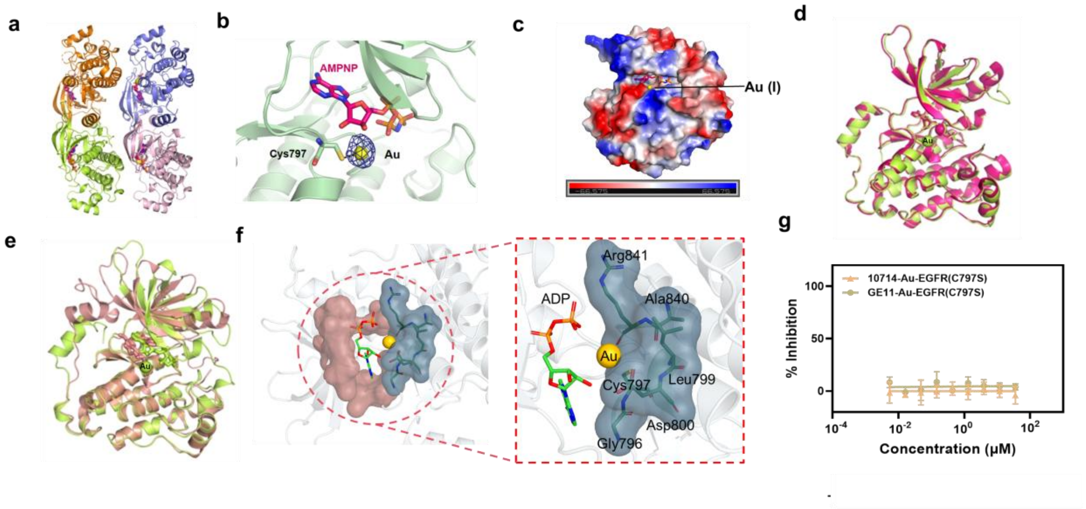


**Figure S12. Structural and functional analysis of Au binding to Cys797 of EGFR T790M. (a)** Overall structure of the EGFR–Au complex within one crystallographic asymmetric unit. The four EGFR molecules are shown in cartoon representation and form two dimers, colored orange, green, blue and pink. AMP-PNP molecules are shown as sticks and Au atoms as yellow spheres. **(b)** Enlarged view of the Au–S interaction at Cys797. The anomalous difference Fourier map for Au is shown as a blue mesh contoured at ~3σ. **(c)** Electrostatic potential surface of the EGFR–Au complex, with positive and negative potentials shown in blue and red, respectively, and Au shown as a gold sphere. **(d)** Structural comparison of Au-bound EGFR T790M (residues 696–1022; green) with the corresponding unbound structure (red; PDB 4ZSE). Au is shown as a green sphere. The Mg ion associated with AMP-PNP in the unbound structure (red sphere) was not observed in the Au-bound structure. **(e)** Structural comparison of Au-bound EGFR T790M (green) with osimertinib-bound EGFR T790M (salmon; PDB 6JX4). **(f)** Overall structure illustrating the Au-coordination environment adjacent to the nucleotide-binding pocket of EGFR T790M. The Au-coordination region surrounding Cys797 is highlighted in blue and the remainder of the nucleotide-binding pocket in salmon. The enlarged view shows the region comprising Gly796, Cys797, Leu799, Asp800, Ala840 and Arg841. **(g)** Enzymatic activity of EGFR C797S following incubation with 10714–Au or GE11–Au.

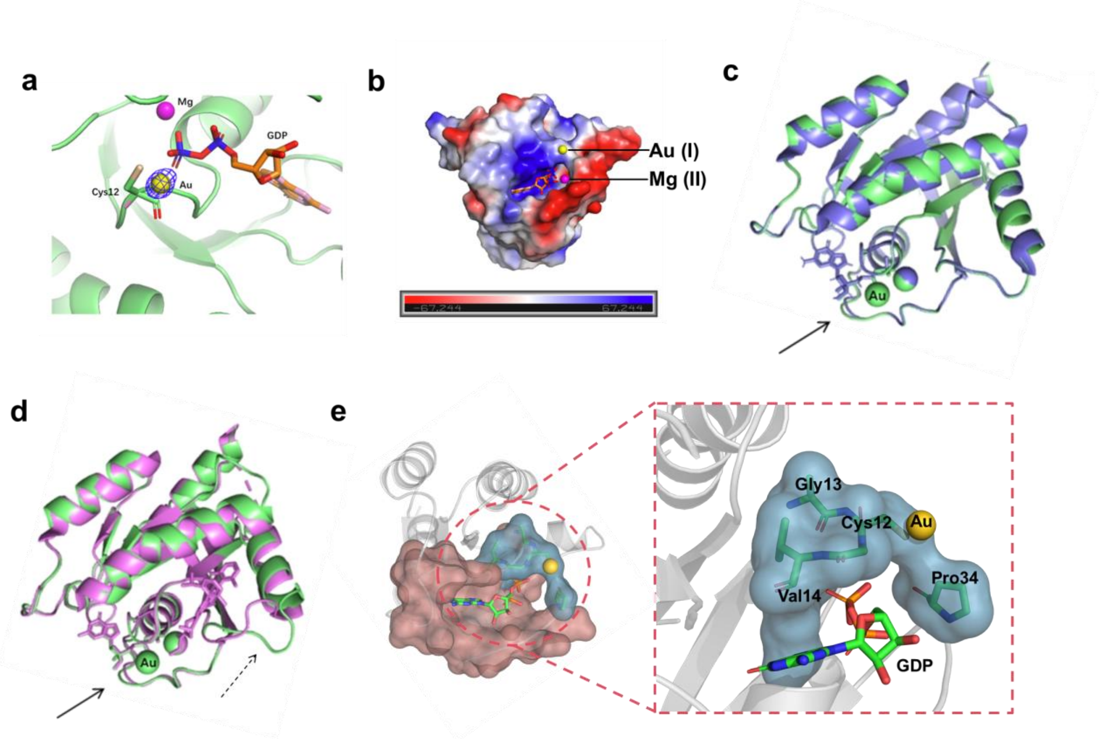


**Figure S13. Structural analysis of Au binding to Cys12 of KRAS G12C. (a)** Enlarged view of the Au–S interaction at Cys12. The anomalous difference Fourier map for Au is shown as a blue mesh contoured at ~5σ. **(b)** Electrostatic potential surface of the KRAS G12C–Au complex, with positive and negative potentials shown in blue and red, respectively. **(c)** Structural comparison of Au-bound KRAS G12C (green) with a previously reported GDP-bound KRAS G12C structure (blue; PDB 4LDJ). The arrow indicates the conformational difference in the Switch I region. **(d)** Structural comparison of Au-bound KRAS G12C (green) with KRAS G12C covalently bound to AMG 510 (purple; PDB 6OIM). The solid and dashed arrows indicate conformational differences in the Switch I and Switch II regions, respectively. **(e)** Overall structure illustrating the Au-coordination environment adjacent to the GDP-binding pocket of KRAS G12C. The Au-coordination region surrounding Cys12 is highlighted in blue and the remainder of the GDP-binding pocket in salmon. The enlarged view shows the region comprising Cys12, Gly13, Val14 and Pro34.

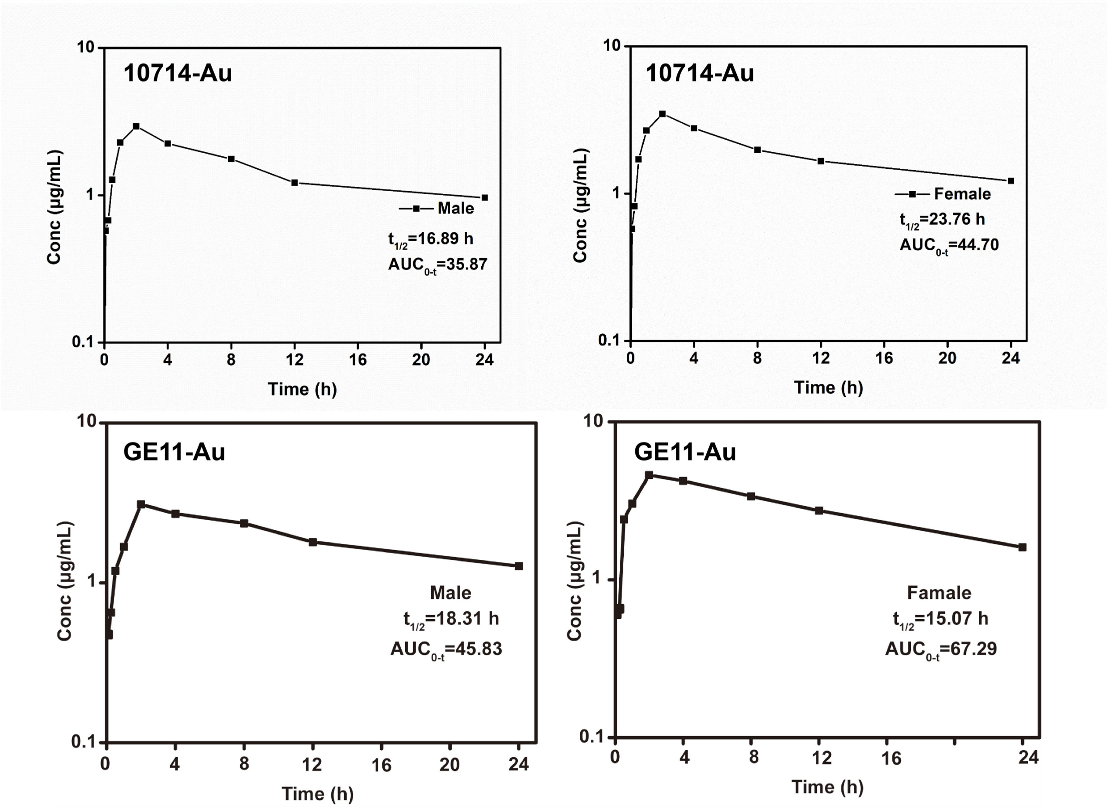


**Figure S14. Pharmacokinetics of 10714–Au and GE11–Au in male and female rats.** Plasma Au concentration–time profiles following administration of the indicated Au-delivery molecules.

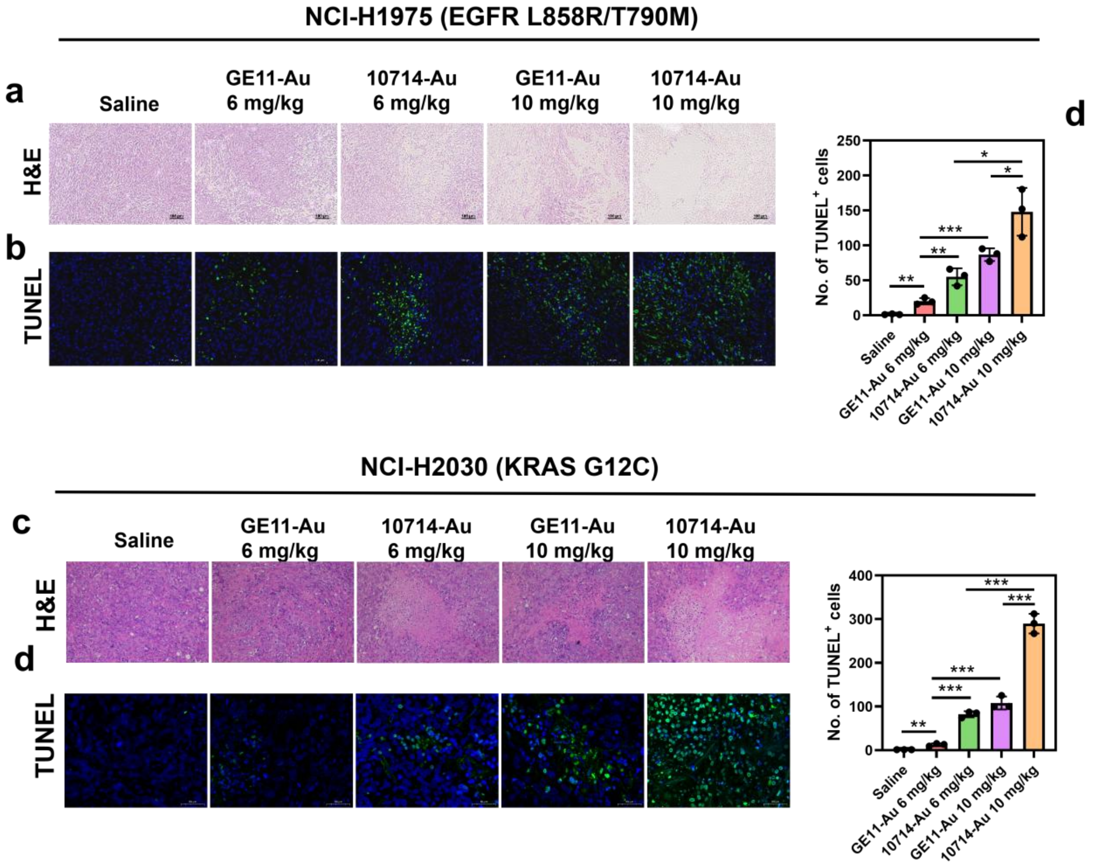


**Figure S15. Effects of Au-delivery molecules on tumor cell death in NSCLC xenograft models.** **(a)** H&E staining of NCI-H1975 tumor tissue following the indicated treatments. Scale bar, 100 μm. **(b)** TUNEL staining and quantification of NCI-H1975 tumor tissue following the indicated treatments. Scale bar, 50 μm. **(c)** H&E staining of NCI-H2030 tumor tissue following the indicated treatments. Scale bar, 100 μm. **(d)** TUNEL staining and quantification of NCI-H2030 tumor tissue following the indicated treatments. Scale bar, 50 μm.

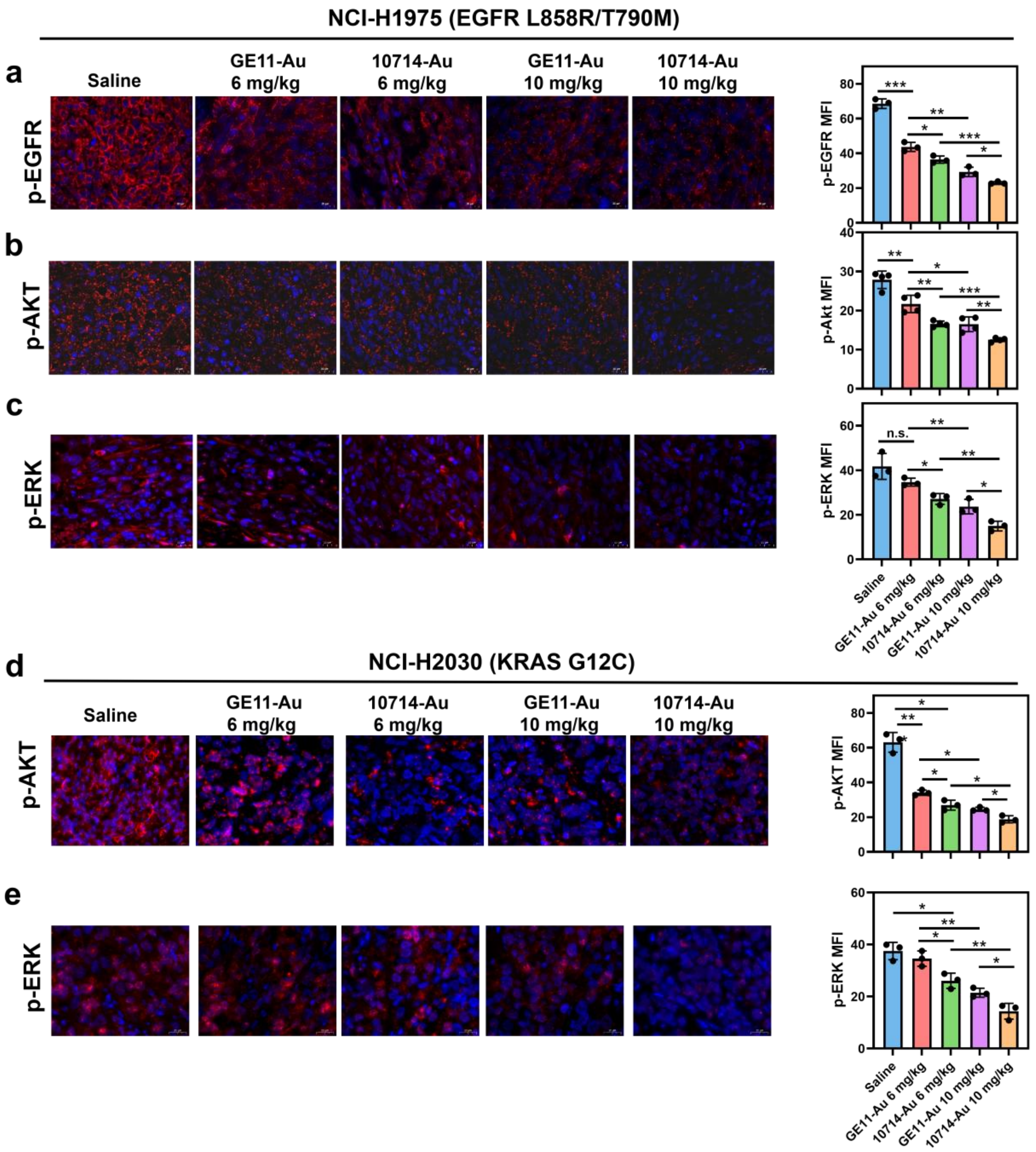


**Figure S16. Au-delivery molecules suppress EGFR and KRAS downstream signals in xenograft cancer models. (a-c)** Immunofluorescence staining of p-EGFR (a), p-AKT (b), and p-ERK (c) on NCI-H1975 tumor tissue and the statistics calculated by Image J was shown. Scale bar=20 μm. **(d, e)** Immunofluorescence staining of p-AKT (d) and p-ERK (e) on NCI-H2030 tumor tissue and the statistics was shown. Scale bar=20 μm.

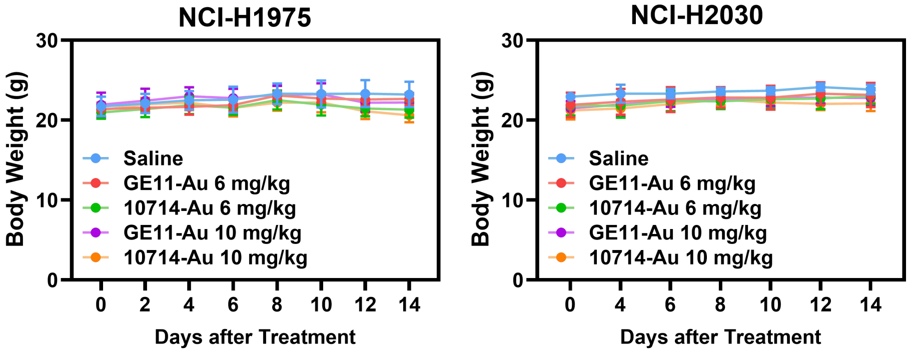


**Figure S17. Body weight during treatment of NSCLC xenograft-bearing mice.** Body weight was monitored for 14 days in mice bearing the indicated xenografts and receiving the indicated treatments.

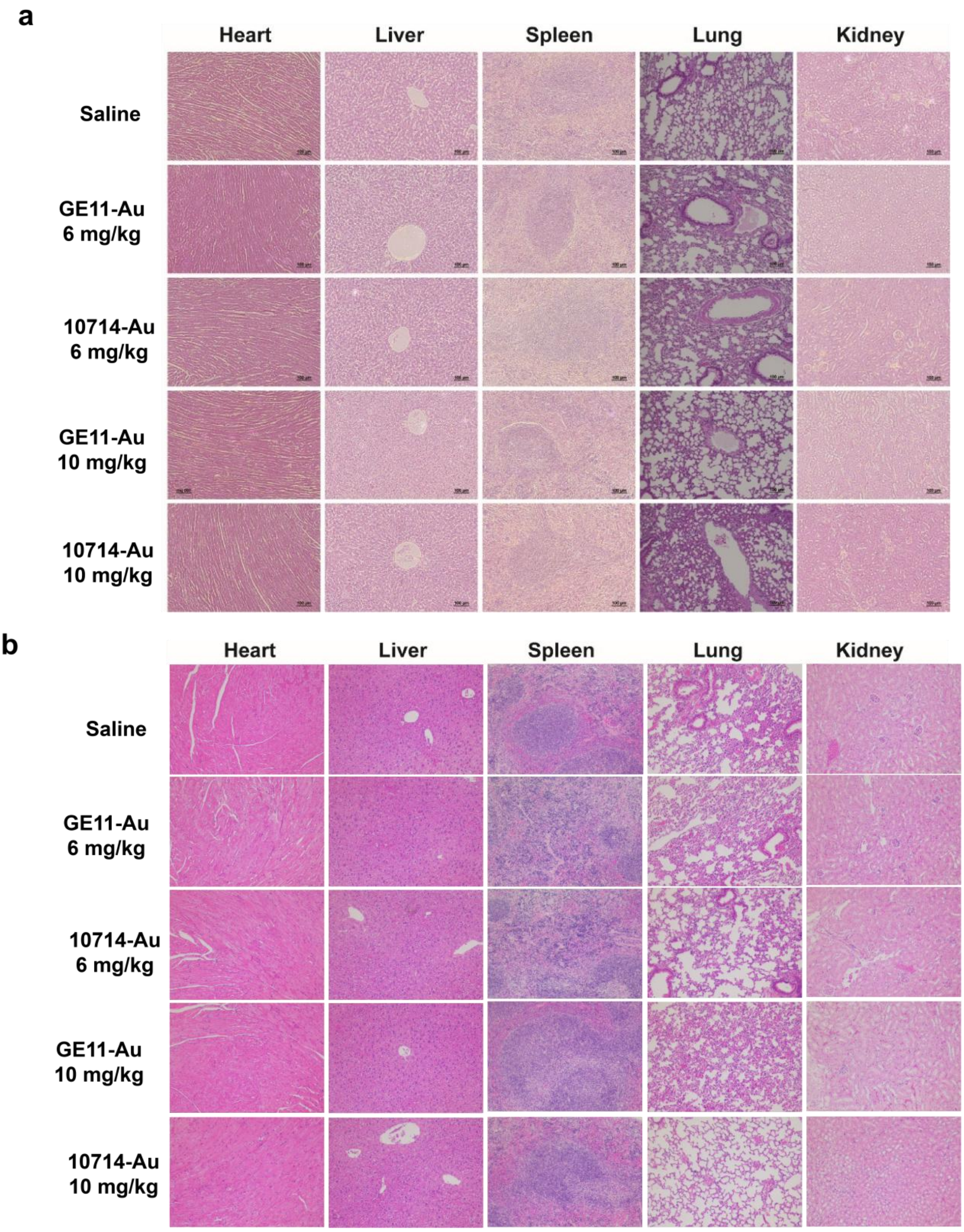


**Figure S18. Histological analysis of major organs following treatment of NSCLC xenograft-bearing mice. (a,b)** H&E staining of major organs from mice bearing (a) H1975 or (b) H2030 xenografts after 14 days of the indicated treatments. Scale bar, 100 µm.

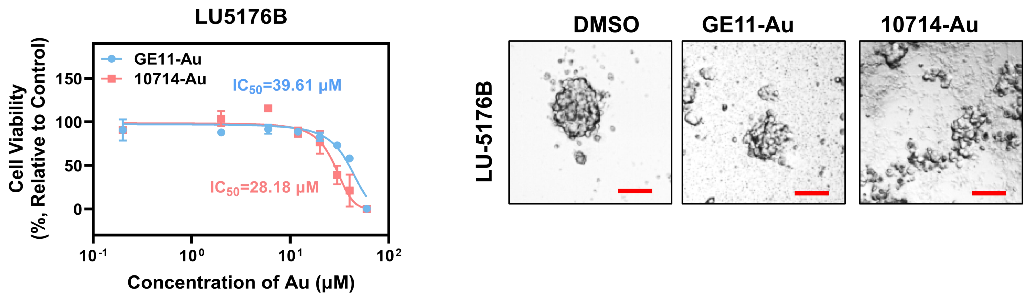


**Figure S19. Effects of 10714–Au and GE11–Au in a third KRAS G12C patient-derived organoid model.** LU5176B organoids were treated with the indicated concentrations of 10714–Au or GE11–Au. Cell viability and representative images following treatment are shown. Scale bar, 100 μm.

### Supplementary Tables

**Table S1. EGFR target experimental peptides collected from the literature.**

| ID | Peptide sequence |
|---|---|
| 1 | YHWYGYTPQNVI |
| 2 | WQTNYIHPYVYG |
| 3 | YGPWYNHYITQ |
| 4 | AHWYGYTPQNV |
| 5 | YAWYGYTPQNV |
| 6 | YHAYGYTPQNV |
| 7 | YHWAGYTPQNV |
| 8 | YHWYAYTPQNV |
| 9 | YHWYGATPQNV |
| 10 | YHWYGYAPQNV |
| 11 | YHWYGYTAQNV |
| 12 | YHWYGYTPANV |
| 13 | YHWYGYTPQAV |
| 14 | YHWYGYTPQNA |
| 15 | YHWYGYTPQN |
| 16 | YHWYGYTPQN |
| 17 | HWYGYTPQNVI |
| 18 | WYGYTPQNV |
| 19 | YHWYGYTPENV |
| 20 | YHWYGYTPQDVI |
| 21 | YKWYGYTPQNV |
| 22 | YRWYGYTPQNV |
| 23 | YHWYGYTPKNV |
| 24 | FHWYGYTPQNV |
| 25 | YHWFGYTPQNV |
| 26 | YHWYGFTPQNV |
| 27 | VVDRVVINNEHQYY |
| 28 | SVEAPIRYYSQTK |
| 29 | RTGEWHYNSKHEL |
| 30 | ENWQLITYGTIL |
| 31 | REWQLQRYGTHW |
| 32 | TDDKIWQQAYGPY |
| 33 | YEYELDEWIRAYGP |
| 34 | LDDIIWQSAYGPTS |
| 35 | WLTYQGGVHFKQHQ |
| 36 | GRYPNVHNLHD |
| 37 | ASFPEELDEWYVAY |

| | |
|---|---|
| 38 | NVEKPLRYWRDTI |
| 39 | VNDIIWQHAYGPN |
| 40 | VNEKVAEVSAGHYS |
| 41 | YANVAIRHTPLPA |
| 42 | DVEHPIRYWAD |
| 43 | LYCDLRKRYIDW |
| 44 | DYKIYNPNSIHNTFD |
| 45 | AAWQLKTYGLVL |
| 46 | TRPPETWRELNI |
| 47 | RQWQLATYGQVY |
| 48 | PEWQIVTYGTVLP |
| 49 | HAYLQNSGGTYW |
| 50 | YDNVRRRLYPL |
| 51 (GE11) | YHWYGYTPQNVI |
| 52 | WQTNYIHPYVYG |
| 53 | YGPWYNHYITQ |
| 54 | AHWYGYTPQNV |

**Table S2. Non-bonded interaction energy of peptide and EGFR.**

| Peptides ID | Non-bonded interaction energy (kcal/mol) |
|---|---|
| 10714 | -43.54 |
| 1648 | -39.98 |
| 697 | -38.86 |
| 4959 | -35.42 |
| 3469 | -33.9 |
| 792 | -33.85 |
| 4163 | -33.21 |
| 2845 | -32.42 |
| 4769 | -31.18 |
| GE11 | -30.51 |
| 3093 | -29.11 |
| 7462 | -28.9 |
| 7891 | -26.74 |
| 4191 | -26.31 |
| 7891 | -26.19 |
| 4191 | -26.13 |
| 7339 | -22.16 |
| 6150 | -21.71 |
| 10176 | -21.18 |
| 2417 | -18.37 |
| 5694 | -17.9 |
| 9360 | -17.54 |
| 7967 | -17.48 |

**Table S3. Zdock and ADCP dock simulations of the selected 5 peptides compared to GE11.**

| Peptides ID | Zdock | ADCP |
|---|---|---|
| 697 | 1735.973 | -53.538378 |
| 3469 | 1749.597 | -52.887051 |
| 10714 | 1972.862 | -44.299891 |
| 4959 | 1727.033 | -42.948435 |
| 1648 | 1721.197 | -42.82844 |
| GE11 | 1549.433 | -11.305573 |

**Table S4. Candidate peptides for synthesis**

| ID | Peptide sequence |
|---|---|
| 697 | RWNQLHRYYAH |
| 1648 | VEWQLKRYGFIL |
| 3469 | ANEAERRYGDWY |
| 4959 | WAEYIDEWGYYY |
| 10714 | KVWRLRKYRLE |
| GE11 | YHWYGYTPQNVI |

**Table S5. X-ray diffraction data collection, refinement and validation statistics of indicated complexes**

| | **EGFR-Au(I)** | **KRAS-Au(I)** | **KRAS-Native** |
|---|---|---|---|
| **Data collection** | | | |
| Diffraction source | SSRF-BL02U1 | SSRF-BL10U2 | SSRF-BL10U2 |
| Detector | Pilatus3 S 6M | Eiger X 16M | Eiger X 16M |
| Wavelength (Å) | 0.98 | 0.98 | 0.98 |
| Space group | *P*21 | *P*212121 | *P*212121 |
| **Cell dimensions** | | | |
| a, b, c (Å) | 70.9, 101.0, 87.3 | 39.1, 41.0, 91.7 | 39.0, 40.2, 91.1 |
| α, β, γ (°) | 90, 102.2, 90 | 90, 90, 90 | 90, 90, 90 |
| Resolution (Å) | 69-3.1 (3.3-3.1)[a] | 50-1.3 (1.35-1.30)[a] | 50-1.75 (1.81-1.75)[a] |
| *Rmerge* | 0.153 (0.654) | 0.073 (0.282) | 0.056(0.187) |
| *Rpim* | 0.076 (0.318) | 0.023 (0.218) | 0.018 (0.124) |
| *I/σI* | 8.1 (2.8) | 19.1 (3.1) | 23.1 (4.5) |
| Completeness (%) | 94.4 (99.9) | 94.9 (68.5) | 87.5 (81.2) |
| Redundancy | 5.8 (6.0) | 8.9 (2.3) | 8.4 (3.1) |
| Overall B factor from Wilson plot (Å2) | 64.7 | 8.0 | 19.9 |
| **Refinement** | | | |
| Resolution (Å) | 60-3.1 | 50-1.3 | 50-1.75 |
| *Rwork/Rfree* | 0.213/0.257 | 0.169/0.189 | 0.177/0.232 |
| No. reflections | 19,943 | 35,185 | 13,446 |
| **No. atoms** | | | |
| Protein | 9,689 | 1366 | 1366 |
| Ligand/ion | 4/4 | 1/2 | 1/1 |
| Water | 47 | 314 | 141 |
| B-factor | 60.7 | 13.5 | 24.7 |
| **R.m.s deviations** | | | |
| Bond lengths (Å) | 0.005 | 0.005 | 0.007 |
| Bond angles (°) | 0.8 | 0.9 | 1.0 |
| **Poor rotamers (%)** | 0 | 0 | 0 |
| **Ramachandran plot** | | | |
| Favored (%) | 96.9 | 98.8 | 98.8 |
| Allowed (%) | 2.9 | 1.2 | 1.2 |
| Disallowed (%) | 0.2 | 0 | 0 |
| **PDB code** | | | |

**Table S6-1. The direct interactions between Au(I) and EGFR/AMP-PNP with the distances less than 4 Å**

| | **EGFR [Atom name]** | **Distance [Å]** |
|---|---|---|
| **Au(I) ion** | Arg 841 [O] | 3.8 |
| | Cys 797 [SG] | 2.3 |
| | Asp 800 [OD1] | 3.9 |
| **Au(I) ion** | **AMP-PNP [Atom name]** | **Distance [Å]** |
| | AMP-PNP [O2'] | 3.6 |
| | AMP-PNP [O3'] | 3.8 |

**Table S6-2. The direct interactions between Au(I) and EGFR/ADP with the distances less than 4 Å**

| | EGFR [Atom name] | Distance [Å] |
|---|---|---|
| **Au(I) ion** | Arg 841 [O] | 3.8 |
| | Cys 797 [SG] | 2.3 |
| | Asp 800 [OD1] | 3.9 |
| **Au(I) ion** | **ADP [Atom name]** | **Distance [Å]** |
| | ADP [O2'] | 3.6 |
| | ADP [O3'] | 3.9 |

**Table S7. The direct interactions between Au(I) and KRAS/GDP with the distances less than 5 Å**

| | KRAS [Atom name] | Distance [Å] |
|---|---|---|
| **Au(I) ion** | Gly 13 [N] | 3.7 |
| | Cys 12 [O] | 3.8 |
| | Cys 12 [SG] | 2.2 |
| **Au(I) ion** | **GDP [Atom name]** | **Distance [Å]** |
| | GDP [O2B] | 5.0 |
| **Au(I) ion** | **H2O [Atom name]** | **Distance [Å]** |
| | HOH 7 [O] | 4.0 |
| | HOH 161 [O] | 2.7 |

**Table S8. Hematological analysis of H2030 tumor-bearing mice after 14 days of indicated treatments**

| Analyte | Unit | Saline | GE11-Au 6 mg/kg | 10714-Au 6 mg/kg | GE11-Au 10 mg/kg | 10714-Au 10 mg/kg | Normal parameter range |
|---|---|---|---|---|---|---|---|
| WBC | $10^9$/L | 6.48±0.97 | 6.32±0.61 | 6.05±0.67 | 6.19±0.69 | 6.16±0.50 | 0.80-10.60 |
| LYM# | $10^9$/L | 4.35±0.49 | 4.35±0.33 | 4.03±0.16 | 4.87±0.13 | 4.34±0.58 | 0.60-8.90 |
| MONO# | $10^9$/L | 0.31±0.02 | 0.30±0.02 | 0.30±0.01 | 0.30±0.02 | 0.30±0.02 | 0.04-1.40 |
| NEUT# | $10^9$/L | 1.87±0.09 | 1.90±0.15 | 1.85±0.10 | 1.97±0.17 | 1.84±0.12 | 0.23-3.60 |
| EOS# | $10^9$/L | 0.09±0.02 | 0.09±0.01 | 0.08±0.03 | 0.09±0.02 | 0.08±0.03 | 0.00-0.51 |
| BASO# | $10^9$/L | 0.05±0.01 | 0.04±0.01 | 0.04±0.02 | 0.04±0.01 | 0.04±0.01 | 0.00-0.12 |
| LYM% | % | 58.4±4.28 | 57.5±4.08 | 57.83±2.72 | 59.63±3.41 | 57.33±3.62 | 40.0-92.0 |
| MONO% | % | 3.43±0.23 | 3.87±0.21 | 3.53±0.21 | 3.77±0.15 | 3.67±0.15 | 0.9-18.0 |
| NEUT% | % | 34.17±4.63 | 38.53±2.25 | 36.30±2.52 | 38.87±1.43 | 36.5±2.31 | 6.5-50.0 |
| EOS% | % | 1.53±0.25 | 1.67±0.21 | 1.70±0.10 | 1.53±0.31 | 1.57±0.21 | 0.0-7.5 |
| BASO% | % | 0.53±0.12 | 0.50±0.20 | 0.47±0.15 | 0.53±0.15 | 050±0.10 | 0.0-1.5 |
| RBC | $10^{12}$/L | 8.70±0.27 | 8.65±0.41 | 8.37±0.15 | 8.70±0.51 | 8.32±0.34 | 6.5-11.5 |
| HGB | g/L | 133.00±4.58 | 137.00±4.00 | 139.00±2.65 | 135.00±3.00 | 131.00±2.00 | 110-165 |
| HCT | % | 42.00±1.97 | 42.00±2.65 | 43.20±1.40 | 44.80±1.54 | 41.93±1.56 | 35.0-55.0 |
| MCV | fL | 44.57±1.58 | 46.73±1.94 | 46.63±2.08 | 46.40±2.55 | 45.97±1.96 | 41.0-55.0 |
| MCH | pg | 16.33±0.95 | 16.43±1.23 | 16.5±0.61 | 15.60±0.70 | 15.47±0.59 | 13.0-18.0 |
| MCHC | g/L | 334.00±6.24 | 335.00±12.49 | 335.67±6.11 | 337.67±4.16 | 334.00±11.79 | 300-360 |
| RDW-SD | fL | 33.20±0.36 | 33.73±2.51 | 33.50±1.06 | 31.10±1.44 | 33.47±0.70 | 23.0-39.0 |
| RDW-CV | % | 15.47±0.78 | 15.43±0.91 | 15.40±0.75 | 14.83±0.35 | 15.73±0.40 | 12.0-19.0 |
| PLT | $10^9$/L | 1111.67±12.66 | 1154.67±47.59 | 1101.33±22.12 | 1101.67±25.70 | 1061.33±72.15 | 400-1600 |
| MPV | fL | 5.33±0.15 | 5.30±0.20 | 5.37±0.25 | 5.33±0.15 | 5.23±0.42 | 4.0-6.2 |
| PCT | % | 0.64±0.04 | 0.61±0.01 | 0.61±0.03 | 0.67±0.02 | 0.63±0.03 | 0.100-0.780 |
| PDW | | 16.33±0.55 | 15.83±0.45 | 16.07±0.15 | 16.17±0.23 | 15.93±0.25 | 12.0-17.5 |

**Table S9. Blood biochemical analysis of H2030 tumor-bearing mice after 14 days of indicated treatments**

| Analyte | Unit | Saline | GE11-Au 6 mg/kg | 10714-Au 6 mg/kg | GE11-Au 10 mg/kg | 10714-Au 10 mg/kg | Normal parameter range |
|---|---|---|---|---|---|---|---|
| GLU | mmol/L | 5.36±0.5 | 5.02±0.11 | 4.95±0.09 | 5.08±0.13 | 5.62±0.41 | 4.66-13.42 |
| ALT | U/L | 49.83±2.84 | 50.87±3.85 | 49.3±3.75 | 52.58±2.84 | 49.26±4.02 | 10.06-96.47 |
| AST | U/L | 169.89±8.67 | 171.62±12.48 | 169.77±12.69 | 173.57±8.82 | 165.15±8.36 | 36.31-235.48 |
| ALB | g/L | 28.73±1.27 | 29.18±1.84 | 28.37±1.36 | 29.67±1.42 | 29.02±1.74 | 21.22-39.15 |
| TG | mmol/L | 1.69±0.07 | 1.72±0.12 | 1.69±0.09 | 1.76±0.08 | 1.69±0.12 | 0.84-2.72 |
| LDL-C | mmol/L | 0.17±0.02 | 0.18±0.02 | 0.17±0.02 | 0.18±0.02 | 0.17±0.02 | 0.12-0.26 |
| HDL-C | mmol/L | 1.48±0.04 | 1.52±0.08 | 1.47±0.06 | 1.53±0.05 | 1.49±0.06 | 1.28-2.65 |
| CREA | μmol/L | 24.81±1.85 | 25.24±2.65 | 24.21±2.68 | 26.49±1.81 | 23.75±1.5 | 10.91-85.09 |
| UREA | mg/dL | 11.9±0.82 | 12.14±1.09 | 11.93±1.3 | 12.25±0.97 | 11.82±1.19 | 10.81-34.74 |
| CK | U/L | 798.24±16.32 | 812.83±17.82 | 799.25±29.52 | 808.7±13.76 | 801.63±24.01 | 0-2070.55 |
| UA | μmol/L | 139.22±11.14 | 142.91±22 | 131.92±23.69 | 143.54±12.13 | 132.43±15.35 | 44.42-224.77 |
| LDH | U/L | 507.23±64.67 | 512.18±35.94 | 538.21±27.18 | 519.55±44.35 | 499.59±37.97 | 157.41-899.72 |

**Table S10. The EGFR and KRAS mutation status in Organoid**

| Sample Name | EGFR mutation | KRAS mutation |
|---|---|---|
| KO-29091 | T790M | WT |
| KO-65534 | T790M | WT |
| LU11692B | p.R521K | p.G12C |
| LU11786B | WT | p.G12C |
| LU5176B | p.R521K | p.G12C |
